# Disease-linked mutations in hnRNPA2B1 accelerate condensate maturation and promote amyloid aggregation

**DOI:** 10.64898/2026.08.28.747731

**Authors:** Luis Fernando Durán-Armenta, Rani Van der Eecken, Annelore Stroobants, Khadija Wahni, Joris Messens, Dominique Maes, Remy Loris, Peter Tompa

## Abstract

Heterogeneous nuclear ribonucleoprotein A2B1 (hnRNPA2B1) is a multifunctional RNA-binding protein that undergoes liquid-liquid phase separation (LLPS), contributing to the assembly of membraneless organelles across different cellular contexts. However, dysregulated phase separation can drive the transition from functional condensates to pathological protein aggregates. Two mutations in the low-complexity domain (LCD) of hnRNPA2B1 (D302V and P310L) have been linked to neurodegenerative diseases. How these mutations alter the interplay between LLPS and aggregation remains poorly understood. Here, we show that these disease-linked mutations accelerate condensate maturation and promote amyloid-like aggregation of hnRNPA2B1, with fibrils emerging from condensate-like cores. To dissect the mechanistic link between phase separation, aggregation, and disease, we focused on a conserved 25-amino-acid region within the LCD harboring both mutation sites. Deletion of this region markedly impairs LLPS and abolishes aggregation, revealing its contribution to both processes. Isolated peptides derived from this region have an intrinsic propensity for amyloid-like fibril formation, which is enhanced by the mutations. Although unable to phase separate, these peptides remodel condensate morphology and nucleate aggregation of the LCD. Together, our results identify a sequence-encoded mechanism linking disease-associated mutations to altered condensate behavior and amyloid aggregation, providing molecular insights into how aberrant phase transitions may contribute to hnRNPA2B1-associated disease.

**Significance Statement:** We address how disease-linked mutations in hnRNPA2B1 alter the interplay between phase separation and pathological aggregation. We demonstrate that the D302V and P310L mutations accelerate condensate maturation and promote fibril formation. By dissecting its low-complexity domain (LCD), we functionally characterized a short amyloidogenic region that contributes to phase separation and is critical for aggregation. Peptides derived from this region form amyloid fibrils that promote ThT-positive assembly and morphological remodeling of the LCD, with disease-linked mutations enhancing this response. Our findings establish an intrinsic sequence-encoded contribution linking disease-linked mutations to accelerated condensate maturation and amyloid aggregation, providing a mechanistic baseline for future studies incorporating physiological RNA partners.

## Introduction

Heterogeneous nuclear ribonucleoproteins A2 and B1 (hnRNPA2 and hnRNPB1) are two RNA-binding proteins produced by alternative splicing of the *HNRNPA2B1* gene. hnRNPA2 is the predominant isoform, accounting for ∼ 90% of total *HNRNPA2B1* expression in most human tissues (1). Each isoform consists of two folded RNA recognition motifs (RRMs) and a C-terminal intrinsically disordered low-complexity domain (LCD). These proteins differ only by an N-terminal 12-residue-long segment (positions 3-14), encoded by exon 2, which is present exclusively in hnRNPB1 (1, 2). Because functional differences between the two isoforms remain poorly characterized (3, 4), hnRNPA2 and hnRNPB1 are often collectively referred to as hnRNPA2B1 in the literature. For consistency, we use hnRNPA2B1 to denote the B1 isoform throughout this study and number all the residues according to its sequence, including those in the isolated LCD and its derived peptides.

hnRNPA2B1 plays essential roles in RNA metabolism, including transcriptional regulation (5), alternative splicing (6, 7), mRNA stability and decay (8, 9), mRNA transport (10), and translational control (5, 7). Beyond these functions, hnRNPA2B1 is a core component of several membraneless organelles (MLOs), including stress granules (SGs) (11), RNA transport granules (12), and hnRNP particles (13). MLOs, also referred to as biomolecular condensates, form through liquid-liquid phase separation (LLPS), a process driven by multivalent interactions among proteins, nucleic acids, and other cellular components (14, 15).

The phase separation behavior of hnRNPA2B1 is driven by its LCD, which mediates interactions with other hnRNPA2B1 molecules, nucleic acids (16) and other phase-separating proteins (11, 17–19). The LCD also governs the material properties and the long-term stability of the resulting condensates. Within this domain, specific short-sequence motifs known as low-complexity, amyloid-like, reversible kinked segments (LARKS) adopt kinked β-strands that assemble through backbone hydrogen bonding into labile β-sheets, promoting reversible self-association and LLPS (20–22). In contrast, the steric zipper motif GNYNDF forms β-sheets that pack tightly through self-complementary interfaces, giving rise to highly stable amyloid fibrils (22–24).

The balance between reversible phase separation and stable amyloid fibrils is reflected in the maturation of hnRNPA2B1 condensates into more solid, gel-like states (25, 26). Although condensates are normally disassembled once they are no longer needed, their reversibility can be disrupted by aging, persistent cellular stress, or pathogenic mutations, promoting an irreversible transition toward solid aggregates, which are often implicated in disease (27). For instance, hnRNPA2B1, together with other SG-associated proteins, has been identified in cytoplasmic inclusions in neurodegenerative diseases such as amyotrophic lateral sclerosis (ALS) and multisystem proteinopathy (MSP) (23, 27, 28).

Two mutations within the LCD of hnRNPA2B1, D302V and P310L (D290V and P298L in hnRNPA2) have been linked to MSP (23) and Paget’s disease of bone (29), respectively. The D302V mutation affects the aspartic acid residue within the steric zipper motif, enhancing amyloid fibril formation (23, 30–32). Cryo-EM structures of mCherry-hnRNPA2 LCD fibrils have revealed that this mutation causes the PY-nuclear localization signal (PY-NLS) to become buried within the fibril core, potentially explaining the cytoplasmic accumulation of hnRNPA2B1 in affected tissues (33). In contrast, the P310L variant has been studied much less extensively. Computational models suggest that this mutation may reduce fibril stability and that its pathogenicity instead may instead arise from impaired nuclear import (32).

Only few studies have addressed the effect of the disease-linked mutations on the LLPS behavior of hnRNPA2B1 (26, 33), the vast majority primarily focusing on protein aggregation. However, growing evidence suggests that pathological aggregation may represent an endpoint of a broader and more complex process driven by aberrant phase separation (34, 35). Understanding how these mutations alter early phase separation, condensate stability, and aging is therefore essential for elucidating the molecular mechanisms underlying disease.

In this study, we present a systematic biophysical characterization of full-length wild-type and mutant hnRNPA2B1 variants and their isolated LCDs under near-native conditions (36) to determine how these mutations alter condensate formation, maturation, and accelerate the progression toward amyloid aggregation. To probe the mechanistic link between LLPS and aggregation, we focused on a previously identified amyloidogenic region within the fibril core (22, 33), which encompasses the steric zipper motif and the two disease-associated mutation sites. Using isolated peptides and a deletion construct, we determined how this region contributes to phase separation, structural ordering, and amyloid formation. By integrating studies on the full-length protein, its isolated LCD, and derived peptides, we establish a multiscale mechanistic framework describing how specific sequence features and disease-linked mutations modulate condensate formation, maturation, and amyloid aggregation.

## Results

### Disease-linked mutations accelerate hnRNPA2B1 condensate maturation

Previous studies have shown that full-length hnRNPA2B1 (26, 37) and its isolated LCD (16, 26, 33, 36) can undergo phase separation. However, phase behavior is highly sensitive to storage conditions and sample preparation, with dilution from strong denaturants (26, 33) or cleavage of solubility tags (26) leading to different kinetic outcomes (36). Here, we induce LLPS by diluting proteins from a high-pH storage buffer into a zwitterionic buffer at physiological pH, allowing condensate formation to be monitored at near-native conditions without residual denaturants or solubility tags (36, 37).

To assess the effect of the disease-linked mutations on condensate formation and maturation, we systematically compared the phase behavior of full-length wild-type hnRNPA2B1 against the D302V and P310L mutants. As shown by turbidity measurements, all three hnRNPA2B1 variants readily undergo phase separation, with no detectable differences in their kinetics **[Fig. 1A]**. Thus, disease-linked mutations do not measurably affect initial condensate formation under these conditions. When visualized under the microscope, the newly formed spherical condensates readily bind thioflavin T (ThT), suggesting that structural ordering occurs at very early stages of condensate formation **[Fig. 1B]**. However, ThT can also bind to folded proteins, and each RRM consists of a four-stranded antiparallel β-sheet with two α-helices (38). To exclude the possibility that the observed ThT fluorescence originated from interactions with the RRMs, we assessed ThT binding with the isolated LCD constructs. Condensates formed by all three LCD variants also exhibit ThT fluorescence **[*SI Appendix*, Fig. S1A]**, supporting the presence of structural ordering within the LCD.

**Fig 1.**
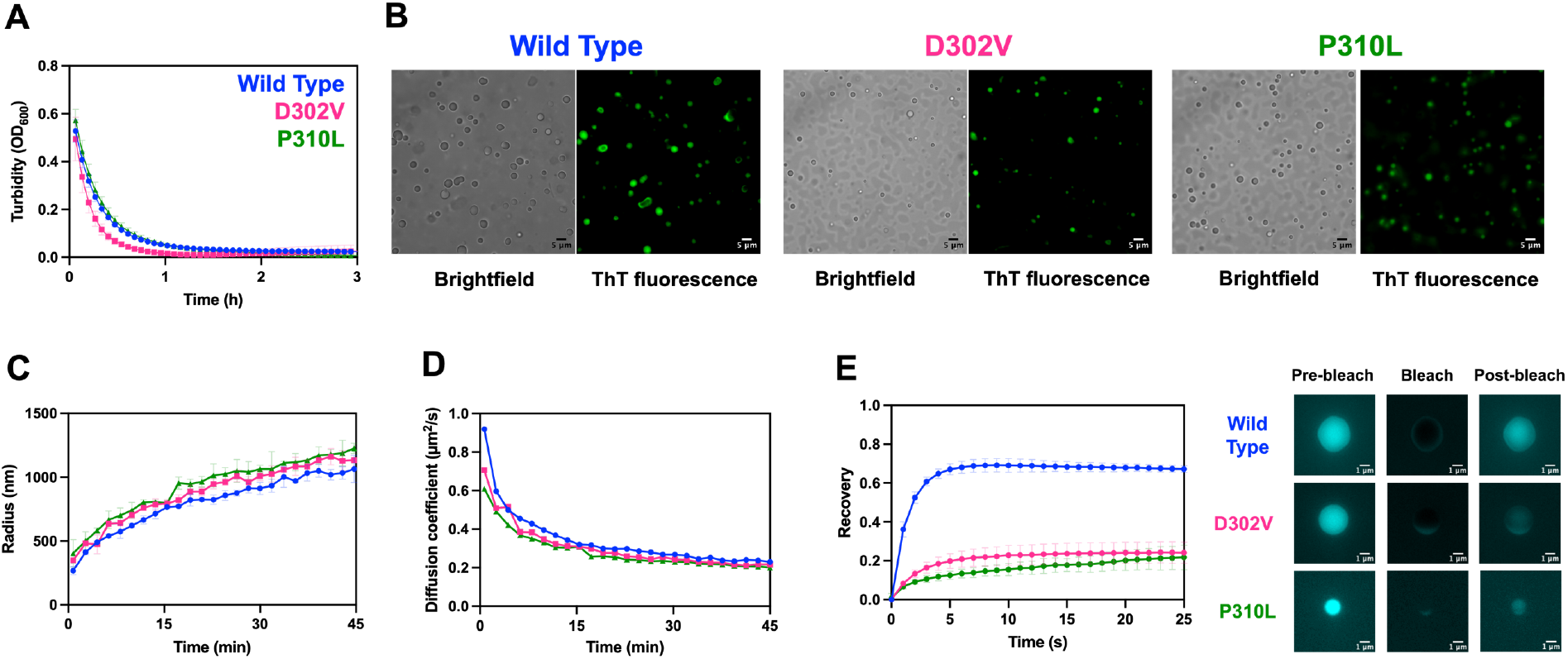
Disease-linked mutations alter the material properties of hnRNPA2B1 condensates. **A)** Turbidity measurements (OD_600_) of wild type (blue), D302V (pink), and P310L (green) hnRNPA2B1 following LLPS induction. **B)** Brightfield and thioflavin T (ThT) fluorescence microscopy of newly formed hnRNPA2B1 condensates. Scale bar: 5 µM. **C)** Condensate growth kinetics followed by evolution of the average hydrodynamic radius (nm) determined by DLS. **D)** Time-dependent diffusion coefficients (µm^2^/s) calculated from DLS measurements. **E)** Fluorescence recovery after photobleaching (FRAP) recovery curves of hnRNPA2B1 condensates (left) and representative pre-bleach, bleach, and 25 s post-bleach images (right). Scale bar: 1 µM. Data are presented as mean ± SD. Turbidity measurements (A) represent three technical replicates. DLS measurements (C and D) correspond to three independent experiments. FRAP recovery curves (E) represent five individual condensates.

Next, we monitored condensate formation and maturation by dynamic light scattering (DLS). Consistent with our previous findings (37), the earliest detectable wild-type hnRNPA2B1 condensates display initial average hydrodynamic radius ∼ 300 nm and grow over time following the characteristic t^1/3^ scaling expected for diffusion-driven coarsening (39). Under the same conditions, the disease-linked mutants, D302V and P310L, display slightly larger condensates (∼ 350 nm and ∼ 400 nm, respectively) that exhibit faster growth kinetics and lower diffusion coefficients (*D*_*t*_), consistent with accelerated condensate coarsening and altered material properties **[Fig. 1C, D, *SI Appendix*, Fig. S2]**. To directly assess condensate viscoelasticity, we performed fluorescence recovery after photobleaching (FRAP) assays. WT condensates recover ∼ 70% of their fluorescence within seconds, indicating liquid-like behavior. In contrast, D302V and P310L exhibit markedly reduced recovery (∼ 25%) over the same period [**Fig. 1E]**. Together, the DLS and FRAP measurements indicate that the disease-linked mutations accelerate condensate maturation from highly dynamic liquid-like states toward gel-like states. Similar differences in condensate growth and molecular mobility are observed for the isolated LCD constructs **[*SI Appendix*, Figure S1]**. However, these effects were more pronounced in the full-length proteins, indicating that the consequences of the disease-linked mutations cannot be fully attributed to the intrinsic properties of the LCD but are also influenced by the molecular context of the full-length protein.

The accelerated maturation of disease-linked variants raised the possibility that the mutations promote higher order oligomerization of hnRNPA2B1 before condensate formation. Mass photometry of soluble D302V and P310L yield apparent molecular masses of 73 ± 12 kDa and 68 ± 2 kDa, respectively **[SI Appendix, Fig. S3]**. These values are comparable to the previously determined molecular mass for WT hnRNPA2B1 (79 ± 8 kDa) (37), indicating that both mutants retain the predominantly dimeric state of the wild-type protein. Therefore, the accelerated condensate maturation is not caused by differences in the initial oligomeric state.

### Disease-linked mutations accelerate the transition from condensates to amyloid fibrils

Given their accelerated condensate maturation, we next investigated whether D302V and P310L also accelerate the transition toward amyloid-like aggregation. To this end, aggregation was monitored by ThT fluorescence following LLPS induction and complemented with endpoint fluorescence microscopy. Wild type hnRNPA2B1 displays a characteristic sigmoidal increase in ThT fluorescence following a lag phase **[Fig. 2A]**. This rate-limiting phase is shortened for both disease-linked mutants, resulting in an earlier rise and substantially higher ThT fluorescence. These effects are particularly predominant for the P310L mutant, which exhibits the highest aggregation propensity. Endpoint fluorescence microscopy reveals distinct aggregate morphologies for each variant **[Fig. 2B, *SI Appendix*, Fig.S4A]**. WT hnRNPA2B1 forms interconnected networks of ThT-positive assemblies, in which protein-rich regions are linked by thin filamentous structures. In contrast, D302V forms larger, irregular ThT-positive assemblies interconnected by prominent filamentous extensions, whereas P310L forms extensive fibrillar-like networks. The effect of the disease-linked mutations was even more pronounced in the isolated LCD variants. Both D302V and P310L display shorter lag phases and strongly accelerated aggregation kinetics compared to WT LCD, with P310L exhibiting the strongest effect **[Fig. 2C]**. Striking differences in aggregate morphology were observed after overnight incubation **[Fig. 2D, *SI Appendix*, Fig.S4B]**. WT LCD predominantly forms large rounded ThT-positive species. In contrast, D302V and P310L form irregular, condensate-like, protein-rich assemblies from which fibril-like structures extend. These filamentous networks are much denser in the P310L mutant. Together, these findings demonstrate that the disease-linked mutations consistently accelerate the transition toward amyloid-like assemblies in both the full-length hnRNPA2B1 and the isolated LCD, supporting a mechanistic link between condensate maturation and fibrillization.

**Fig 2.**
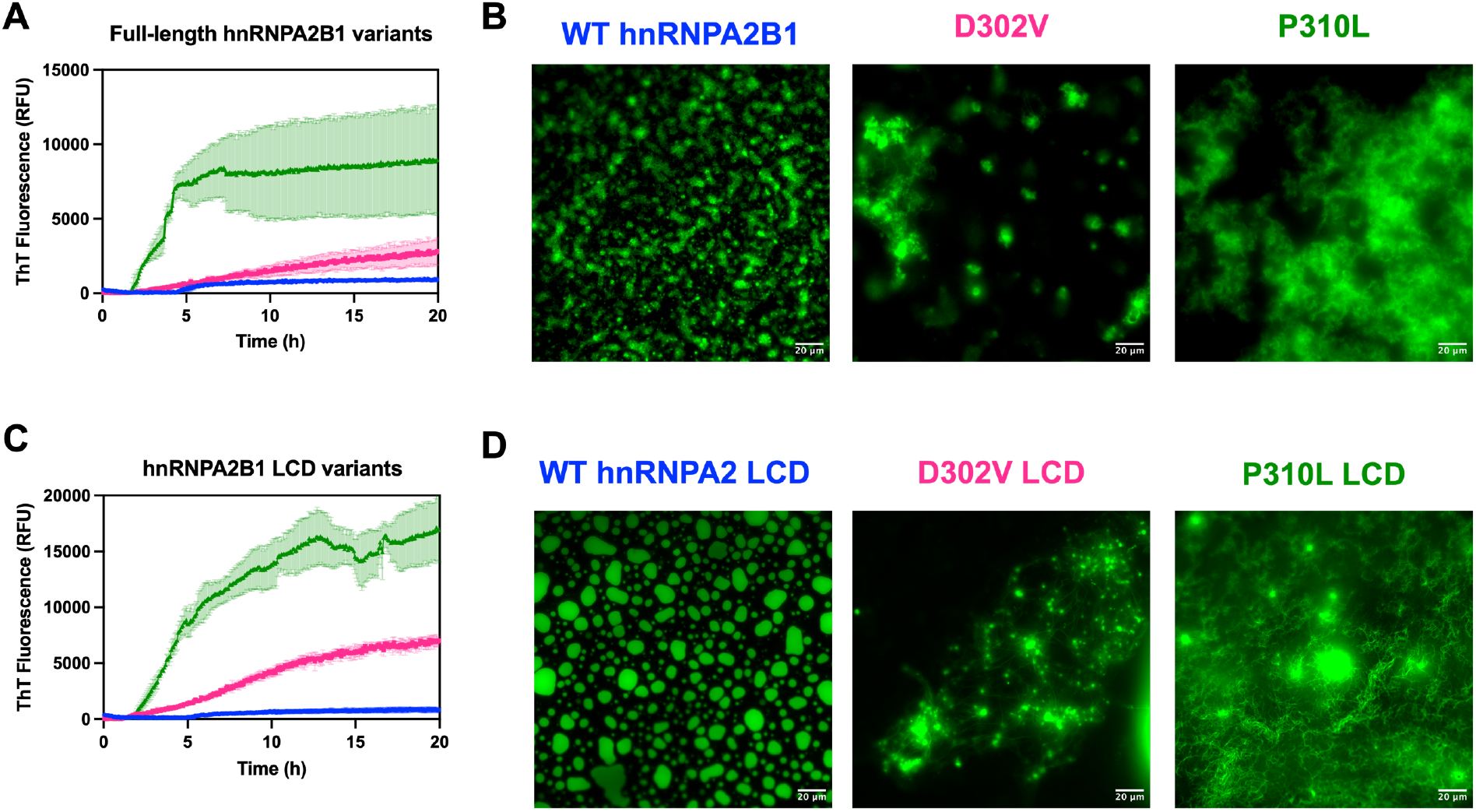
Disease-linked mutations enhance amyloid-like aggregation of hnRNPA2B1 and its LCD. **A)** ThT fluorescence kinetics of WT, D302V, and P310L full-length hnRNPA2B1 following induction of phase separation. **B)** Representative fluorescence images of ThT-stained full-length hnRNPA2B1 variants after 20 h incubation. **C)** ThT fluorescence kinetics of WT, D302, and P310L hnRNPA2B1 LCD. **D)** Representative fluorescence micrographs of ThT-stained hnRNPA2B1 LCD variants after 20 h incubation. Data are presented as the mean ± SD of three technical replicates. Scale bars: 20 µm.

### Disease-linked mutations reside within an evolutionarily conserved amyloidogenic region

To investigate the molecular basis of the accelerated condensate maturation and aggregation observed in the disease-linked variants, we focused on the residues surrounding D302 and P310 and examined their evolutionary conservation and amyloidogenic propensity. Both D302 and P310 are highly conserved across 392 vertebrate hnRNPA2B1 LCD orthologs, while the disease-linked mutations are strongly disfavored in their respective positions, reflecting evolutionary pressure against introducing bulky hydrophobic residues. Notably, many of the surrounding residues are likewise strongly conserved, indicating that the D302 and P310 are part of a broader sequence element. **[*SI Appendix*, Fig.S5A]**. We then used ZipperDB (40) to examine how these mutations affect the amyloidogenic propensity of the surrounding sequence. Consistent with previous reports, the D302V mutation enhances the previously identified steric zipper (23, 26, 30), whereas P310L extends it (26) **[*SI Appendix*, Fig.S5B, Table S1]**.

Together, these analyses define a conserved, mutation-sensitive segment spanning G291-P315 of hnRNPA2B1 (G279-P303 in hnRNPA2), hereafter referred to as the Amy region, that overlaps with the fibril cores previously identified in mCherry-hnRNPA2 LCD structures (22, 33) **[Fig. 3A]**.

**Fig 3.**
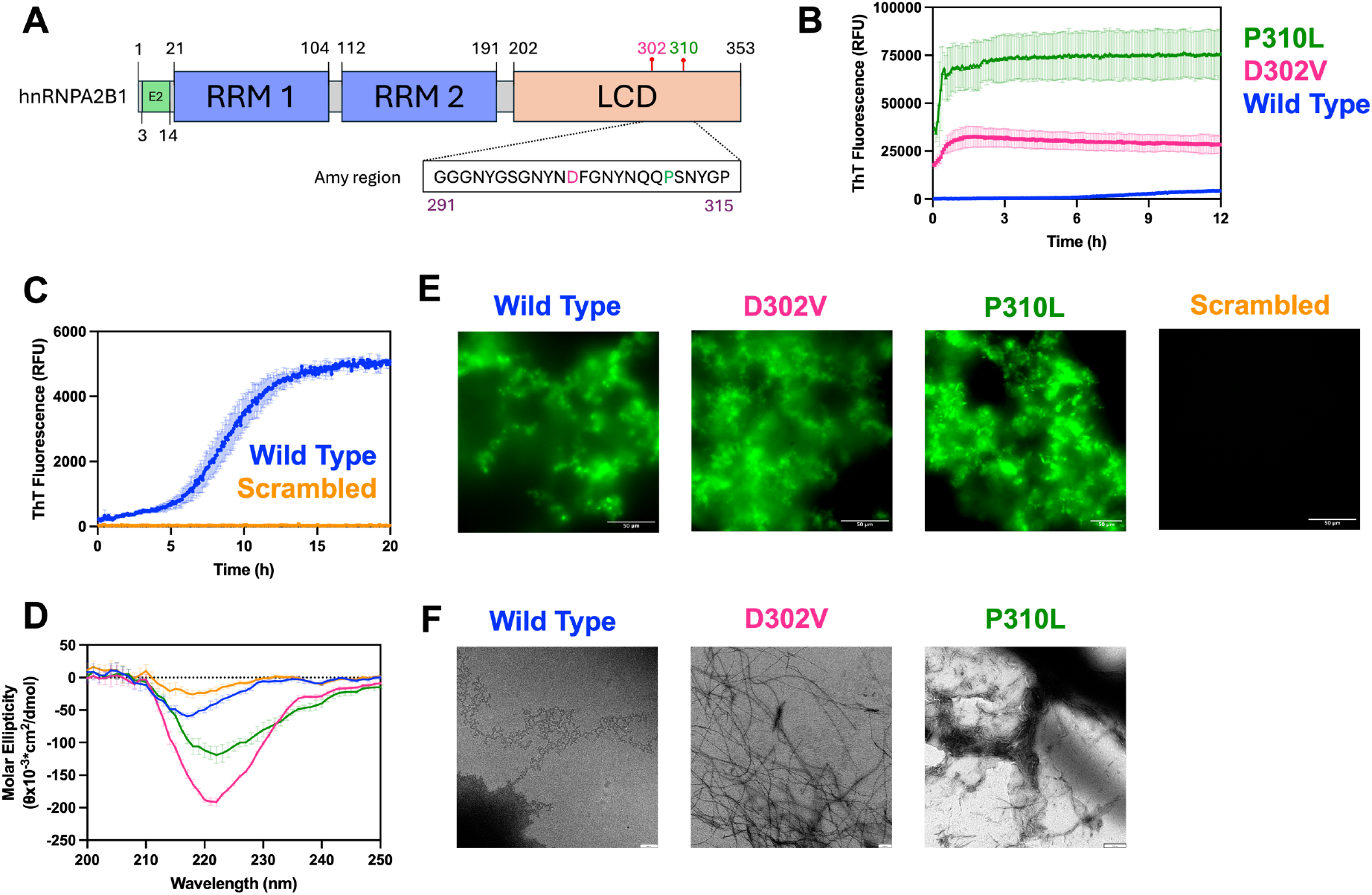
The Amy region within the LCD of hnRNPA2B1 is intrinsically amyloidogenic and its aggregation is enhanced by disease-linked mutations. **A)** Schematic representation of the domain architecture of hnRNPA2B1: two globular RNA-recognition motifs (RRMs; blue) and a disordered low-complexity domain (LCD; orange). The B1 isoform has an additional N-terminal segment encoded by exon 2 (E2; green box). The disease-linked mutations D302V (pink) and P302L (green) are indicated by the red pins. The location of the Amy region (G291-P315) is indicated by the inset. **B)** ThT fluorescence kinetics of wild-type (blue), D302V (pink), and P310L (green) peptides. **C)** ThT fluorescence kinetics of WT (blue) and scrambled (orange) peptides. Data are presented as the mean ± SD of three technical replicates. **D)** Circular dichroism spectra of WT (blue), D302V (pink), P310L (green), and scrambled (orange) recorded after the pH jump. Data are presented as the mean ± SD of three independent experiments. Representative **E)** fluorescence microscopy images and **F)** negative-stain transmission electron micrographs of isolated Amy region peptides following aggregation. The scrambled control showed no detectable ThT fluorescence and did not form fibrils. Scale bars: 50 μm for ThT fluorescence microscopy, 200 nm for electron microscopy.

### Disease-linked mutations enhance the intrinsic amyloidogenicity of the Amy region

To experimentally examine whether the Amy region is intrinsically amyloidogenic, we synthesized peptides corresponding to the WT, D302V, and P310L sequences together with a scrambled control (SC), generated by randomizing the native sequence order **[*SI Appendix* Fig. S6]**. We first examined the phase separation behavior of the Amy peptides across concentrations ranging from 10 μM to 75 μM. None of the peptides exhibit detectable LLPS, as determined by turbidity measurements **[*SI Appendix* Fig. S7]**, indicating that the Amy region alone is insufficient to drive phase separation, and that additional multivalent interactions distributed along the LCD are required. Despite the absence of LLPS, the WT and mutant Amy peptides display sigmoidal ThT fluorescence kinetics. Both D302V and P310L exhibit substantially higher ThT fluorescence than the WT peptide from the first time point, followed by a rapid increase that reaches a plateau within the first hours [**Fig. 3B**]. Across the tested conditions, the P310L variant consistently yields the highest ThT signal, followed by D302V, and lastly by WT **[*SI Appendix* Fig. S8]**, mirroring the increased aggregation propensity observed for the full-length protein and isolated LCDs. To further determine whether the observed aggregation was sequence-specific, the WT peptide was compared against the scrambled control. While WT exhibits a characteristic sigmoidal increase in ThT fluorescence, the scrambled peptide remains at baseline throughout the recording [**Fig. 3C**], indicating that aggregation strongly depends on the native sequence order rather than on amino acid composition alone.

We next examined the secondary structure propensity of 0.2 mg/mL (∼75 μM) peptide solutions by CD spectroscopy. Upon dilution from the storage buffer to physiological pH, the WT, D302V, and P310L peptides adopt β-sheet-like signatures, with pronounced minima around 218-220 nm, whereas the scrambled control shows minimal structural ordering **[*SI Appendix* Fig. S9]**. WT develops a moderate spectrum that remains stable over 3 hours, whereas D302V displays the strongest β-sheet-like features **[Fig 3D, *SI Appendix* Fig. S10]**. P310L similarly exhibits a strong β-sheet-like structure at early timepoints, although its rapid aggregation precludes measurements after longer incubations **[*SI Appendix* Fig. S11]**. In contrast, the scrambled peptide lacks a well-defined β-sheet-like spectrum and shows minimal changes over time.

Finally, fluorescence microscopy reveals extensive interconnected ThT-positive aggregates for WT, D302V, and P310L, whereas no detectable fluorescence was observed for the scrambled control **[Figure 3E]**. Negative-stain transmission electron microscopy confirms that these assemblies correspond to amyloid fibrils, with D302 and particularly P310L forming dense networks of overlapping and intertwined fibrils **[Figure 3F]**. Together, these results demonstrate that the Amy region possesses an intrinsic, sequence-specific propensity to adopt β-sheet-like conformations and assemble into amyloid fibrils under physiological conditions. Disease-linked mutations further enhance this amyloidogenicity by accelerating structural ordering and promoting extensive fibril formation.

### The Amy region modulates phase separation and aggregation of hnRNPA2 LCD

Having established that the isolated Amy region is sufficient to drive amyloid formation, but not phase separation, we next investigated its contribution to these processes within the context of the LCD by generating a deletion variant lacking this region (ΔAmy; ΔG291-P315). Following pH-jump in HEPES buffer, ΔAmy exhibits no detectable phase separation by either turbidity measurements or fluorescence microscopy **[*SI Appendix* Fig. S12A]**. However, ΔAmy retains a reduced capacity to phase separate in phosphate buffer, indicating that the Amy region strongly contributes to LCD-driven condensation, but it is not strictly required for it. ΔAmy condensates show little or no ThT fluorescence **[*SI Appendix* Fig. S12B]**, suggesting that condensates can form without detectable β-sheet-rich assemblies. Moreover, ΔAmy remains ThT-negative after overnight incubations **[*SI Appendix* Fig. 13]**, further confirming the role of the Amy region as a driver of amyloid-like aggregation. Consistent with this behavior, its CD spectrum exhibits low-amplitude without the β-sheet-like signature observed for the isolated Amy peptides **[*SI Appendix* Fig. S14]**, indicating that ΔAmy forms a predominantly disordered ensemble. Together, this identifies the Amy region as a key determinant of hnRNPA2B1 LCD phase behavior. Although other interactions within the LCD can support condensate formation under certain conditions, the Amy region is required for efficient phase separation and is essential for amyloid-like aggregation.

### Amy peptide-derived seeds perturb condensate maturation and promote amyloid aggregation

Finally, we investigated whether preformed fibrils derived from the Amy region could promote amyloid-like aggregation of the LCD through a seeding mechanism. To this end, preformed Amy peptide fibrils were sonicated into smaller fragments and added to wild-type LCD solutions prior to LLPS induction.

Peptide seeding does not affect initial condensate formation, as seeded and unseeded reactions follow similar LLPS kinetics **[Figure 4A]**. Likewise, DLS measurements reveal no major differences in average condensate size or growth kinetics **[Figure 4B, *SI Appendix* Fig. S15]**. Despite these similarities, fluorescence microscopy reveals pronounced differences in condensate morphology **[Figure 4C]**. Unseeded samples and those containing WT or scrambled peptide seeds form spherical condensates. In contrast, disease-linked mutant seeds induce irregular and deformed morphologies. Specifically, the D302V peptide produces a moderate effect, while the P310L peptide results in prominent non-spherical and elongated protein-rich assemblies, suggesting that disease-linked Amy fibrils remodel condensate organization without substantially affecting their formation, growth, or average size. These morphological alterations are accompanied by enhanced amyloid-like aggregation. Preformed Amy fibrils enhance the ThT fluorescence of hnRNPA2B1 LCD, with P310L seeds producing the strongest effect, followed by D302V and WT. In contrast, the scrambled control shows little to no seeding activity **[Figure 4D]**. Together, these findings demonstrate that fibrillar assemblies derived from the Amy region remodel condensate morphology and promote amyloid-like aggregation of the LCD, consistent with a seeding mechanism. Collectively, our experiments demonstrate that the Amy region functions as a sequence-encoded determinant of condensate maturation and amyloid assembly, providing a molecular mechanism by which disease-linked mutations promote pathlogical aggregation.

**Fig 4.**
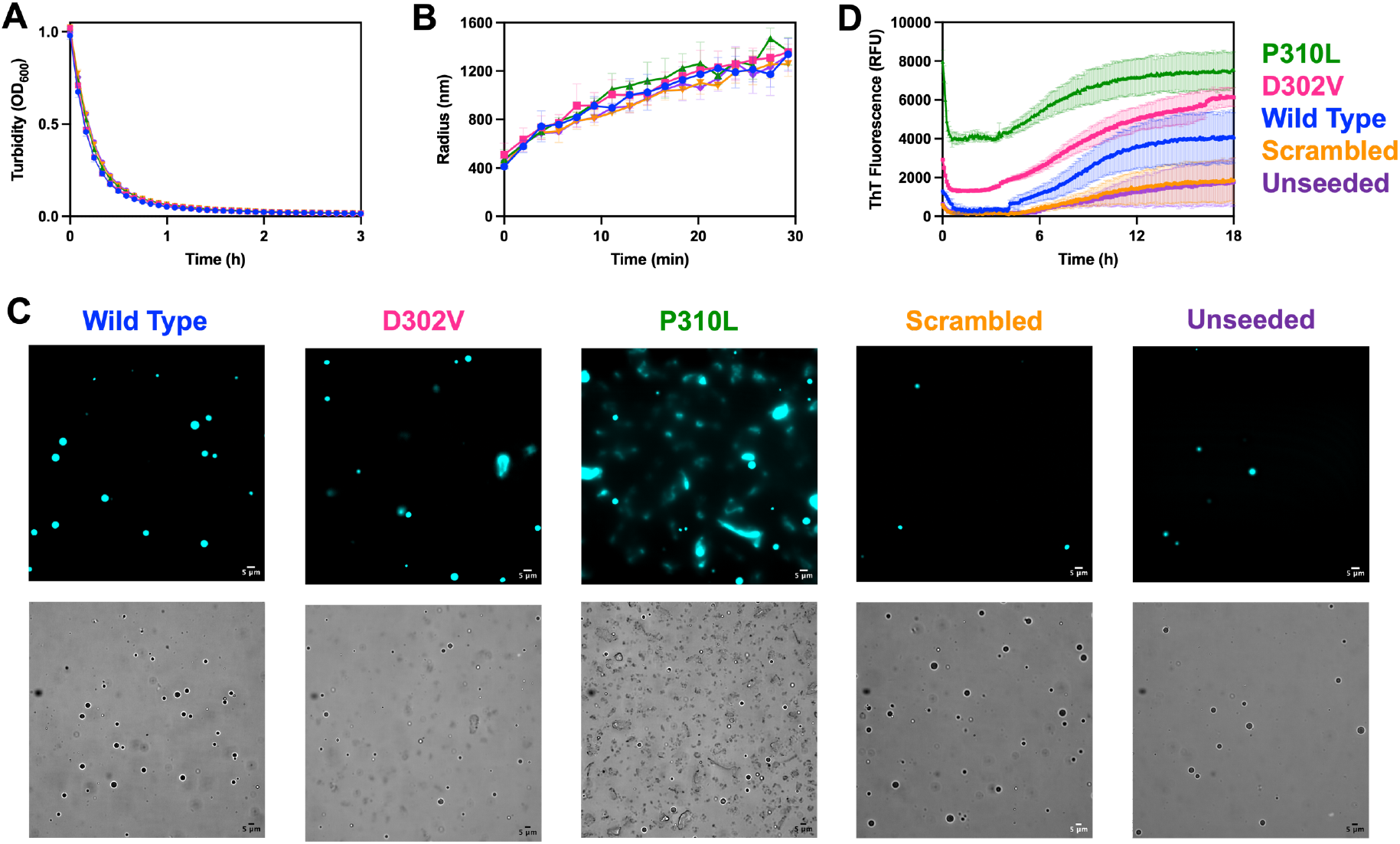
Disease-linked Amy peptide seeds remodel wild type hnRNPA2B1 LCD condensates and nucleate amyloid-like aggregation. **A)** Turbidity measurements of wild type hnRNPA2B1 LCD seeded with preformed WT (blue), D302V (pink), P310L (green), or scrambled (orange) peptide fibrils, or left unseeded (purple). Data plotted represent mean ± SD from three technical replicates. **B)** Seeded and unseeded hnRNPA2B1 LCD condensates. Data plotted represent mean ± SD from three independent experiments. **C)** Representative brightfield and confocal fluorescence microscopy of seeded and unseeded reactions containing DyLight-labeled hnRNPA2B1 LCD. Scale bar: 5μm. **D)** ThT fluorescence kinetics demonstrating enhanced amyloid-like aggregation in reactions seeded with disease-associated peptide fibrils. Data plotted represent mean ± SD from three technical replicates.

## Discussion

Dysregulated phase separation has emerged as central mechanism linking physiological biomolecular condensates to disease-associated protein aggregation. This relationship is strengthened by the enrichment of numerous phase-separating RNA-binding proteins in both stress granules and pathological inclusions (27, 28). Among these proteins, hnRNPA2B1 is associated with multisystem proteinopathy (23) and Paget’s disease of bone (29). Although previous structural and biophysical studies have established that disease-linked mutations accelerate amyloid formation (23, 24, 26, 30, 33), their effects on condensate maturation as well as the molecular mechanisms underlying this transition have remained unclear. Here, we demonstrate that disease-linked mutations accelerate condensate maturation and promote the transition toward amyloid-like aggregation. Furthermore, we functionally establish a conserved region previously identified within the fibril core of mCherry-hnRNPA2 LCD fibrils (22, 33) as the minimal sequence element governing this transition.

Unlike previous studies, in which hnRNPA2B1 phase separation was induced by dilution from denaturants or solubility-tag cleavage (26, 33), our pH-jump approach (36, 37) readily triggers LLPS for both the full-length proteins and the isolated LCDs. Under these conditions, all variants form spherical condensates. These observations highlight both the sensitivity of LLPS to experimental conditions and the versatility of the pH-jump method for investigating phase separation in intrinsically disordered proteins, including those containing folded domains, at near-native conditions.

Our biophysical characterization of full-length hnRNPA2B1 and isolated LCD variants consistently indicates that the disease-linked mutants primarily alter condensate properties, rather than phase separation propensity. Upon formation, wild-type and mutant hnRNPA2B1 condensates readily bind ThT, suggesting that β-sheet-rich structures begin to emerge during the earliest stages of condensate formation, consistent with previous observations of polymeric ordering in liquid-like droplets and hydrogels (20, 21). Although the kinetics of initial phase separation are comparable among the variants, D302V and P310L form larger condensates that mature into less dynamic material states.

Following prolonged incubation, hnRNPA2B1 condensates transition into extensive networks of filamentous, ThT-positive assemblies. Interestingly, both disease-linked mutants form irregular, condensate-like cores from which filamentous structures extend. This morphological progression is accompanied by characteristic sigmoidal ThT aggregation kinetics, both in full-length protein and the isolated LCD. Consistent with previous observations, the D302V mutant shortened the lag phase of amyloid aggregation (30, 33). Interestingly, P310L consistently exhibited the strongest aggregation phenotype across every experimental system examined. To our knowledge, this represents the first biophysical characterization of the aggregation propensity of this disease-linked hnRNPA2B1 variant.

Collectively, our findings place hnRNPA2B1 within an emerging framework in which biomolecular condensates act as intermediates for amyloid aggregation (41–43). Recent studies on the closely related proteins FUS (42) and hnRNPA1 (43) suggest that condensate interfaces promote intermolecular β-sheet formation through local protein density fluctuations, thereby providing favorable sites for amyloid nucleation (42). Rather than representing independent processes, phase separation and amyloid formation therefore appear to constitute part of a broader continuum that is strongly influenced by disease-linked mutations.

While these observations establish that disease-linked mutations accelerate the transition from condensates to amyloid assemblies, they do not explain why D302V and particularly P310L consistently display such strong aggregation phenotypes. Notably, both mutation sites reside within a 54-residue segment previously identified as the core of mCherry-hnRNPA2 LCD fibrils (22, 33). By integrating evolutionary conservation with amyloidogenicity predictions, we refined this region to a highly conserved 25-residue spanning G291-P315 of hnRNPA2B1 (corresponding to G279-P303 in hnRNPA2), hereafter referred to as the Amy region.

Structural characterization of isolated peptides demonstrated that this region possesses an intrinsic propensity to adopt β-sheet-rich conformations and assemble into amyloid fibrils. Although unable to phase separate, the isolated peptides recapitulate the aggregation hierarchy observed in the full-length protein and the LCD models. Whereas previous structural studies indicate that D302V enhances the native steric zipper and stabilizes amyloid fibrillization (23, 26, 30, 32), available predictions suggest that the P310L mutation could reduce fibril stability (32). Our observations instead identify P310L as the most aggregation-prone variant, suggesting that extension of the native steric zipper may generate a longer and more potent amyloidogenic segment than previously appreciated. In contrast, scrambling the native Amy sequence abolishes detectable amyloid assembly, highlighting that the amyloidogenic behavior of this region depends on its native sequence order rather than its amino acid composition alone. Rather than the entire 54-residue fibril core, our results indicate that the 25-residue Amy region is sufficient for the intrinsic amyloidogenic behavior of hnRNPA2B1 LCD.

The intrinsic amyloidogenicity of the isolated Amy peptides raised the question of whether this sequence merely contributes to, or is actually required, for phase separation and amyloid formation within the LCD. The ΔAmy deletion construct is only able to undergo LLPS under the permissive ionic conditions provided by phosphate buffer. The resulting condensates do not display detectable ThT fluorescence and fail to mature into amyloid-like assemblies. Although deletion of the previously reported fibril core abolishes phase separation of hnRNPA2B1 LCD (33), our results demonstrate that removal of only the central 25-residue Amy region partially preserves condensate formation while abolishing detectable amyloid assembly. These observations support a model in which different regions of the LCD mediate phase separation and aggregation (43). Multivalent cation-π interactions between Arg and Tyr residues remain capable of supporting limited condensation in the absence of the Amy region under favorable ionic conditions (43, 44). However, these interactions are not sufficient to drive β-sheet-rich structural transitions required for amyloid assembly. Together, our results identify the Amy region as a central determinant of both condensate formation and their subsequent maturation into amyloid fibrils.

Although previous studies established the self-seeding capacity of hnRNPA2B1 LCD fibrils (23, 24), we sought to determine whether this activity was retained within the isolated 25-residue Amy region. Remarkably, preformed Amy fibrils are sufficient to seed amyloid aggregation of the surrounding LCD, recapitulating the hierarchy observed throughout this study: P310L>D302V>WT. Although the peptide seeds do not exert detectable effect on condensate formation or growth, they profoundly alter condensate morphology. Disease-linked mutant fibril seeds, particularly P310L, induce irregular, elongated condensates with accelerated ThT-positive aggregation, suggesting that pre-formed Amy fibrils remodel condensate maturation rather than initial phase separation.

The large excess of LCD over seeds, the mutation-dependent effects, and the minimal response to scrambled seeds support a model in which Amy fibrils promote templated recruitment of the LCD. Although ThT kinetics and morphological changes support this seeding model, we did not directly track the incorporation of hnRNPA2B1 LCD into the fibrillar material. Nevertheless,these observations remain consistent with emerging evidence that pre-existing amyloid assemblies can actively reshape the material properties of biomolecular condensates. Recent studies have demonstrated that amyloid-rich condensates can transmit their aggregation-prone state to amyloid-poor condensates, thereby propagating pathological material properties (45). The evidence presented here, including evolutionary conservation, amyloidogenicity predictions, isolated peptide behavior, deletion constructs, and peptide-mediated seeding, collectively identifies the Amy region as an important sequence-encoded determinant of LCD-driven amyloid assembly under our experimental conditions.

By using purified proteins and peptides in the absence of RNA, we deliberately isolate the intrinsic, protein-encoded contribution of the LCD and its Amy region to phase-separation and amyloid aggregation. However, our reductionist approach does not capture how RNA and other components of physiological membraneless organelles might modulate the observed mutational hierarchy in condensate maturation and amyloid assembly. Future studies will be needed to determine whether the P310L>D302V>WT hierarchy is maintained in the presence of physiological partners.

Collectively, our findings, provide a mechanistic framework explaining how disease-linked mutations convert physiological condensate maturation into pathological amyloid assembly. The D302V and particularly, the P310L mutations amplify the intrinsic amyloidogenic behavior of a conserved sequence embedded within the LCD of hnRNPA2B1. Through complementary analyses involving the full-length protein, the isolated LCD, minimal peptides, deletion constructs, and seeding assays, we establish the Amy region as the molecular determinant governing the transition from dynamic condensates to amyloid aggregates. Together, our findings support an emerging view in which pathological aggregation represents the endpoint of a broader sequence-encoded assembly pathway initiated by biomolecular condensation and accelerated by disease-linked mutations.

## Materials and Methods

### Evolutionary conservation analysis

Full-length hnRNPA2B1 sequence (UniProt: P22626-1) was used as the query for protein-protein BLAST® with the default parameters. Retrieved homologous sequences were filtered on sequence identity (30%-70%) and query coverage (70%-100%) to generate a multiple sequence alignment (MSA) using Clustal Omega. The RRMs were excluded from the MSA, and a Kullback-Leibler (KL) divergence sequence logo was generated using Seq2Logo (46) with the default parameters.

### Vectors

The pET-22b (+) expression vector encoding 6xHis-SUMO-hnRNPA2B1 (37) was used to generate the D302V and P310L mutants by site-directed mutagenesis. The vectors encoding 6xHis-hnRNPA2 LCD (26) (AddGene #98657) and D290V (16) were kind gifts from Prof. Nicholas L. Fawzi (Brown University, USA) and Dr. Joris van Lindt (VIB-KU Leuven Center for Neuroscience, Belgium), respectively. The P298L mutant and the ΔAmy (ΔG279-P303) construct were generated by site directed mutagenesis. All constructs were verified by Sanger sequencing. LCD constructs were originally generated based on the hnRNPA2 sequence, but this manuscript follows the hnRNPA2B1 residue numbering.

### Protein Expression

All hnRNPA2B1 full-length and LCD variants were expressed in *Escherichia coli* BL21 STAR™ (DE3) cells grown in Terrific Broth (TB) medium as previously described (16, 36, 37). Construct-specific expression conditions are provided in ***SI Appendix***.

### Protein purification

Full-length hnRNPA2B1 variants were purified following our previously established protocol (37). Briefly, the clarified lysate was loaded onto a 5 mL HisTrap™ HP column (Cytiva) equilibrated with wash buffer (50 mM HEPES, 1M NaCl, 10 mM imidazole, 0.5 mM TCEP, pH 8.0). Bound protein was eluted with a linear gradient up to 500 mM imidazole, desalted into Cleavage Buffer (50 mM Tris, 1M NaCl,

0.5 mM TCEP, pH 7.5), and incubated with TEV protease to cleave the 6xHis-SUMO tag. Subsequent purification by reverse HisTrap™ chromatography and size exclusion chromatography (SEC), and dialysis into storage buffer are explained in ***SI Appendix***.

Purified proteins were aliquoted at 15 µM in 10 mM CAPS, pH 11.0, and flash-frozen in liquid nitrogen for storage at -80ºC. Protein identity and purity were verified by intact mass spectrometry and top-down sequencing as previously described (37).

hnRNPA2B1 LCD constructs were purified as described previously (16, 36). Briefly, inclusion bodies were isolated from the cell lysate, solubilized in denaturing buffer (20 mM Tris-Cl, 3 or 8M urea, 500 mM NaCl, 25 mM imidazole, 1 mM DTT, pH 8.0), and purified by immobilized metal affinity chromatography. Bound protein was eluted with 500 mM imidazole, desalted into Cleavage Buffer (50 mM NaH_2_PO_4_, 200 mM NaCl, 3M urea, pH 7.0), and incubated with TEV protease to remove the 6xHis tag. Remaining contaminants were removed by a reverse HisTrap™ chromatography and SEC as explained in ***SI Appendix***. Purified proteins were aliquoted at 50 μM in 10 mM CAPS, pH 11.0, flash-frozen, and stored at -80ºC.

### Peptide design, synthesis, and preparation

Based on the ZipperDB (40) predictions, four peptides encompassing the amyloidogenic region (G291-P315) of hnRNPA2B1 LCD were designed: WT, D302V, P310L, and a scrambled control (SC). The SC sequence was generated using the Shuffle Protein tool of the Sequence Manipulation Suite (Bioinformatics.org) and selected based on low sequence identity (<35%), disruption of the native steric zipper, and amyloidogenicity comparable to the WT peptide. Peptide sequences are described in ***SI Appendix***. All peptides were custom synthesized by GenScript at ≥98% purity. Peptide powders were dissolved in 10 mM CAPS (pH 11.0) to 1.5 mg/mL (≈ 550 μM) and incubated at 55ºC until fully dissolved. Solutions were aliquoted, flash-frozen, and stored at -80 ºC.

### Turbidity measurements

LLPS was induced by diluting the protein stock solutions with 100 mM HEPES, pH 7.5, resulting in final protein concentrations of 20 μM for the hnRNPA2B1 LCD constructs and 10 μM for the full-length hnRNPA2B1 variants. Peptide stock solutions were diluted to the desired final concentrations. Turbidity was recorded at 600 nm every 4 minutes using a Synergy™ HTZ Multi-Mode Microplate Reader (BioTek) at 25 ºC with continuous slow agitation for 3 h. All experiments were performed with at least three technical replicates, and buffer controls were subtracted for baseline correction.

### Dynamic Light Scattering

Dynamic Light Scattering (DLS) measurements were performed using a DynaPro® NanoStar® instrument (Wyatt Technology). Phase-separating protein samples were analyzed at 25ºC by recording scattered light intensity at a fixed angle (95º) using 5 acquisitions of 8 seconds per time point. Data was analyzed and processed as described in ***SI Appendix***. All experiments were performed in triplicate.

### Mass Photometry

Protein landing was recorded using a Refeyn OneMP (Refeyn Ltd) MP system under the same experimental conditions as previously described (37). Briefly, hnRNPA2B1 D302V and P310L were diluted in 10 mM CAPS, pH 11.0 to a final concentration of 75 nM and movies (6,000 frames, 60 s) were acquired. Data was analyzed as explained in ***SI Appendix***. All experiments were performed in triplicate.

### Thioflavin T Fluorescence Measurements

Thioflavin T (ThT) was added to protein and peptide solutions to a final concentration of 25 μM prior to dilution in 100 mM HEPES, pH 7.5. ThT fluorescence was measured every 4 minutes using 450 nm excitation and 490 nm emission filters on the plate reader. Measurements were performed at 25 ºC with slow, continuous agitation for at least 12 hours. All experiments were performed with at least three technical replicates, and buffer fluorescence was subtracted.

### Fluorescent Labeling of Proteins

All hnRNPA2B1 LCD variants were labeled with DyLight® 488 NHS Ester (Thermo Scientific) following the manufacturer’s instructions. Labeling buffer was supplemented with 3M urea to prevent LLPS and aggregation. Unbound dye was removed by extensive dialysis into 10 mM CAPS, pH 11.0. Full-length hnRNPA2B1 was not labeled since this dye artificially increases its aggregation propensity (37).

### Brightfield and Fluorescence Microscopy

Protein and peptide samples were supplemented with the corresponding DyLight ®488 NHS Ester-labeled hnRNPA2B1 LCD variant (1:50) or with ThT (25 μM) prior to the induction of LLPS or aggregation. Brightfield and fluorescence microscopy were performed using an RCM2.5 Re-scan Confocal Microscope system (Confocal.nl; ***SI Appendix***). Images were analyzed using Fiji (ImageJ).

### Fluorescence Recovery After Photobleaching

Fluorescence Recovery After Photobleaching (FRAP) experiments were performed using a Leica DMi8 inverted fluorescence microscope equipped with a FRAP module (Leica Microsystems; ***SI Appendix***). DyLight 488-labeled LCD variants were mixed with an excess of the corresponding unlabeled protein (LCD or full-length hnRNPA2B1) prior to LLPS induction. At least five different stationary condensates were bleached for 500 ms using a 488 nm laser at 100% laser power. Fluorescence intensities were background-corrected and normalized to the mean pre-bleach intensity to generate recovery curves.

### Negative-stain Transmission Electron Microscopy

Peptide samples (0.2 mg/mL) were applied to freshly glow-discharged Formvar carbon-coated copper grids and allowed to adsorb. Grids were subsequently stained with 20 μL droplets of uranyl acetate using a three-step protocol consisting of 10 s in the first droplet, 1 s in the second droplet, and 1 min in the final droplet. Excess stain was removed and the grids were air-dried. Samples were imaged using a JEOL JEM-1400 transmission electron microscope operating at an accelerating voltage of 120 kV.

### Circular Dichroism Spectroscopy

Circular dichroism (CD) spectra were recorded using a BioLogic MOS-500 spectropolarimeter (BioLogic Instruments). Amy peptides and ΔAmy were diluted in 50 mM HEPES (pH 7.0) to a final concentration of 0.2 mg/mL (≈ 75 μM for peptides and ≈20 μM for ΔAmy) and analyzed in a 1 mm quartz cuvette (Hellma Analytics). Far-UV CD spectra (200-250 nm) were recorded at 25ºC using a bandwidth of 1 nm and an acquisition time of 1 s per point. At least five consecutive scans were averaged per sample. Measurements were performed in triplicate using independently prepared samples. Buffer spectra were subtracted, and data was processes as described in ***SI Appendix***.

### Seeding Assays

Amy peptide seeds were generated by incubating WT, D302V, and scrambled peptide solutions (75 μM) overnight at 25 ºC with agitation. Owing to its markedly faster aggregation kinetics, P310L seeds were incubated for only 30 minutes. Fibrils were sonicated (1 min, 1 s on, 1 s off, 20% amplitude) and mixed with WT hnRNPA2B1 LCD to final concentrations of 15 μM (seeds) and 50 μM (LCD) in 100 mM HEPES, pH 7.5. Turbidity and ThT fluorescence were monitored as described above. All experiments were performed with at least three technical replicates, and buffer fluorescence was subtracted.

## Acknowledgements

We sincerely thank Dr. Steven Janvier (VIB-VUB Center for Structural Biology) for intact mass spectrometry and top-down sequencing analyses; Dr. Álvaro Navarro (Fundación Instituto Leloir, Argentina) for generating the multiple sequence alignments used in the conservation analysis; Prof. Tamás Lázár (Structural Biology Brussels) for insightful scientific input and discussions throughout the project, and critical reading of the manuscript; and Axelle Kerstens (VIB Bioimaging Satellite Brussels) for training on confocal microscopy and image processing and analysis. This work was supported by a Fonds Wetenschappelijk Onderzoek (FWO) PhD fellowship (1163625N to L.F.D.A), the Vrije Universiteit Brussel Strategic Research Programs SRP51 and SRP97 (L.F.D.A., D.M., and P.T.), and the European Space Agency (ESA) grant A0-2004-070 (D.M.).

