## Supplementary Appendix for "Disease-linked mutations in hnRNPA2B1 accelerate condensate maturation and promote amyloid aggregation"


**This PDF file includes:**

Supporting text
Figures S1 to S15
Table S1
SI References

#### Supplementary Methods

##### Protein expression

All cultures were grown in Terrific Broth (TB) supplemented with carbenicillin (100 µg/mL) except for the wild type hnRNPA2B1 LCD construct, which was grown in TB supplemented with kanamycin (25 µg/mL). Cultures were incubated at 37ºC with agitation at 180 rpm until an $OD_{600}$ of 0.6–0.8 was reached, after which protein expression was induced with 1 mM IPTG. Full-length hnRNPA2B1 variants and the ΔAmy LCD construct were expressed for 4 hours at 37ºC. All other LCD constructs were expressed overnight at 25ºC. Cells were harvested by centrifugation (5000 g, 15 min, 4ºC).

##### Protein purification of full-length hnRNPA2B1 variants

The cell pellet of 1L of culture was resuspended in 100 mL lysis buffer (50 mM HEPES, 1M NaCl, 10 mM imidazole, 0.5 mM TCEP, pH 8.0) supplemented with 0.1% Triton X, RNAse A, DNAse I, 0.1 mM PMSF, 0.5 mM benzamidine hydrochloride, and 2 cOmplete™ EDTA-free protease inhibitor tablets (Roche). The homogenized solution was sonicated on ice for 10 min (5 s pulse on, 5 s pulse off, 70 % amplitude) and centrifuged (18 000 rpm, 4 ◦C, 1 h). The supernatant was filtered and loaded onto a 5 mL HisTrap™ HP column (Cytiva) equilibrated with wash buffer (50 mM HEPES, 1M NaCl, 10 mM imidazole, 0.5 mM TCEP, pH 8.0). Bound SUMO-hnRNPA2B1 was eluted with a linear gradient of wash buffer supplemented with 500 mM imidazole. The eluate was desalted into Cleavage Buffer (50 mM Tris, 1M NaCl, 0.5 mM TCEP, pH 7.5) using a HiPrep™ 26/10 Desalting column (Cytiva). The solubility tag was cleaved by adding 500 µL of TEV protease (3 mg/mL) and overnight incubation at room temperature. The solution was then loaded onto a second HisTrap™ HP column equilibrated with wash buffer without imidazole. The flow-through containing tag-free hnRNPA2B1 was collected. Remaining contaminants were removed by size exclusion chromatography using a Superdex® 200 Increase 10/300 GL column (Cytiva) equilibrated with 50 mM CAPS, 500 mM NaCl, pH 11.0. Selected fractions were dialyzed into 10 mM CAPS, pH 11.0, aliquoted at 15 µM, and flash-frozen in liquid nitrogen for storage at−80 ºC. Protein concentration was determined using a Nanodrop™ One (Thermo Scientific) using the extinction coefficients (ε) calculated with ProtParam: ε = 41260 $M^{-1}$ $cm^{-1}$ for SUMO-hnRNPA2B1 and ε = 39770 $M^{-1}$ $cm^{-1}$ for hnRNPA2B1. Pure samples showed an A260/A280 ratio of approximately 0.6, indicating minimal nucleic acid contamination.

##### Protein purification of hnRNPA2B1 LCD variants

The cell pellet of 1L of culture was resuspended in 100 mL lysis buffer (20 mM Tris-Cl, 500 mM NaCl, 25 mM imidazole, 1 mM DTT, pH 8.0) supplemented with 0.1% Triton X, RNAse A, DNAse I, 0.1 mM PMSF, 0.5 mM benzamidine hydrochloride, and two cOmplete™ EDTA-free protease inhibitor tablets (Roche). Cells were lysed by sonication for 15 minutes (5 s on / 5 s off, 60% amplitude). Inclusion bodies containing the protein of interest were pelleted by centrifugation (18,000 rpm, 1 h, 4°C). Pellets were resuspended in denaturing buffer (20 mM Tris-Cl, 500 mM NaCl, 25 mM imidazole, 1 mM DTT, pH 8.0) containing either 3M urea (WT and ΔAmy) or 8M urea (D302V and P310L). Inclusion bodies were sonicated again for 10 minutes (10 s on, 10 s off, 70% amplitude) and centrifuged. The lysate was filtered and loaded onto a 5 mL HisTrap™ HP column (Cytiva) equilibrated with denaturing buffer. Bound protein was eluted with a linear imidazole gradient up to 500 mM imidazole. Eluted fractions were desalted into Cleavage Buffer (50 mM $NaH_2PO_4$, 200 mM NaCl, 3M urea, pH 7.0) using a HiPrep™ 26/10 Desalting column (Cytiva) and added 500 µL TEV protease (3 mg/mL). The solution was incubated overnight at room temperature and loaded to a second HisTrap™ HP column equilibrated with denaturing buffer, collecting the flow-through. Remaining contaminants were removed by size-exclusion chromatography on a HiLoad ® 26/600 Superdex ® 200 pg column (Cytiva) equilibrated with 10 mM CAPS, 200 mM NaCl, 3M or 8M urea, pH 11.0. Selected fractions were desalted into 10 mM CAPS, pH 11.0, aliquoted at 50 µM, flash-frozen in liquid nitrogen and stored at -80ºC. Protein concentrations were determined using the theoretical extinction coefficients: ε = 19370 $M^{-1}$ $cm^{-1}$ for ΔAmy and ε = 25330 $M^{-1}$ $cm^{-1}$ for hnRNPA2 LCD, D302V, and P310L. Pure samples showed an A260/A280 ratio of approximately 0.6, indicating minimal nucleic acid contamination.

**Amyloidogenic Peptide Sequences**
**Wild Type**: GGGNYGSGNYNDFGNYNQQPSNYGP
**D302V**: GGGNYGSGNYNVFGNYNQQPSNYGP
**P310L**: GGGNYGSGNYNDFGNYNQQLSNYGP
**Scrambled**: NQNQPGSFGYGYGNYGNNGDYNSPG

**Mass Photometry**
Protein landing was recorded using a Refeyn OneMP (Refeyn Ltd) MP system by diluting 10 µL of hnRNPA2B1 D302V and P310L solutions into a 10 µL drop of filtered 10 mM CAPS, pH 11.0 (Final concentration: 75 nM). Movies (6,000 frames, 60 s) were acquired with the AcquireMP software version 2.1.1 (Refeyn Ltd) using the default settings. Data was analyzed using default settings on DiscoverMP (version 2.1.1; Refeyn Ltd). Contrast-to-mass calibration was performed with MassFerence P1 (Refeyn) using standards of 88, 172, 258, and 344 kDa. The binding and unbinding events were grouped into mass ranges (binning). Frequency distribution with default setting and a bin width of log2(x), where x is the total number of detected particles, was used. Data was represented as the number of particles (counts) vs. mass (kDa). Triplicates were measured, and the average molecular weight and standard deviation were calculated.

**Dynamic Light Scattering**
Data were analyzed using the Dynamics® Software Package (V. 7.10.1.21, Wyatt). The translational diffusion coefficient ($D_t$) was calculated based on the exponential fitting of the autocorrelation function. The hydrodynamic radius was calculated using the Stokes-Einstein equation:

$$D_t = \frac{kT}{6\pi\eta R_h}$$

Where $k$ is the Boltzmann constant, $T$ is the temperature, $\eta$ is the viscosity (assumed to be that of water), and $R_h$ is the measured hydrodynamic radius.

**Circular Dichroism Spectroscopy**
Raw ellipticity values (θ, mdeg) were converted to molar ellipticity [Θ] (degrees*cm$^2$/dmol) using the formula:

$$[\Theta] = \frac{\theta * MW}{10 * L * C}$$

where $\theta$ is ellipticity (mdeg), $MW$ is the molecular weight (g/mol), $C$ is concentration (g/L), and $L$ is path length (cm). Processed spectra were analyzed and exported using BioKine software (BioLogic) and plotted in GraphPad Prism.

**Supplementary information on the microscope setups**

**Brightfield and Fluorescence Microscopy**
Brightfield and fluorescence microscopy were performed using an RCM2.5 Re-scan Confocal Microscope (Confocal.nl) mounted on an Olympus IX3 inverted microscope (Evident Scientific) equipped with an ORCA- Flash 4.0 V3 Digital CMOS camera (Hamamatsu Photonics) and a UPLXAPO 60x/1.42NA oil objective (Olympus). The system was controlled with µManager version 2.0.3.

**Fluorescence Recovery After Photobleaching**
Fluorescence Recovery After Photobleaching (FRAP) experiments were performed using a Leica DMi8 inverted fluorescence microscope equipped with a Leica DFC7000 GT camera, an HC PL FLUOTAR 100x/1.32 Oil PH3 objective, and an integrated FRAP module (Leica Microsystems). The system was controlled using Leica Application Suite X version 3.7.0.20979 (Leica Microsystems).

#### Supplementary Figures

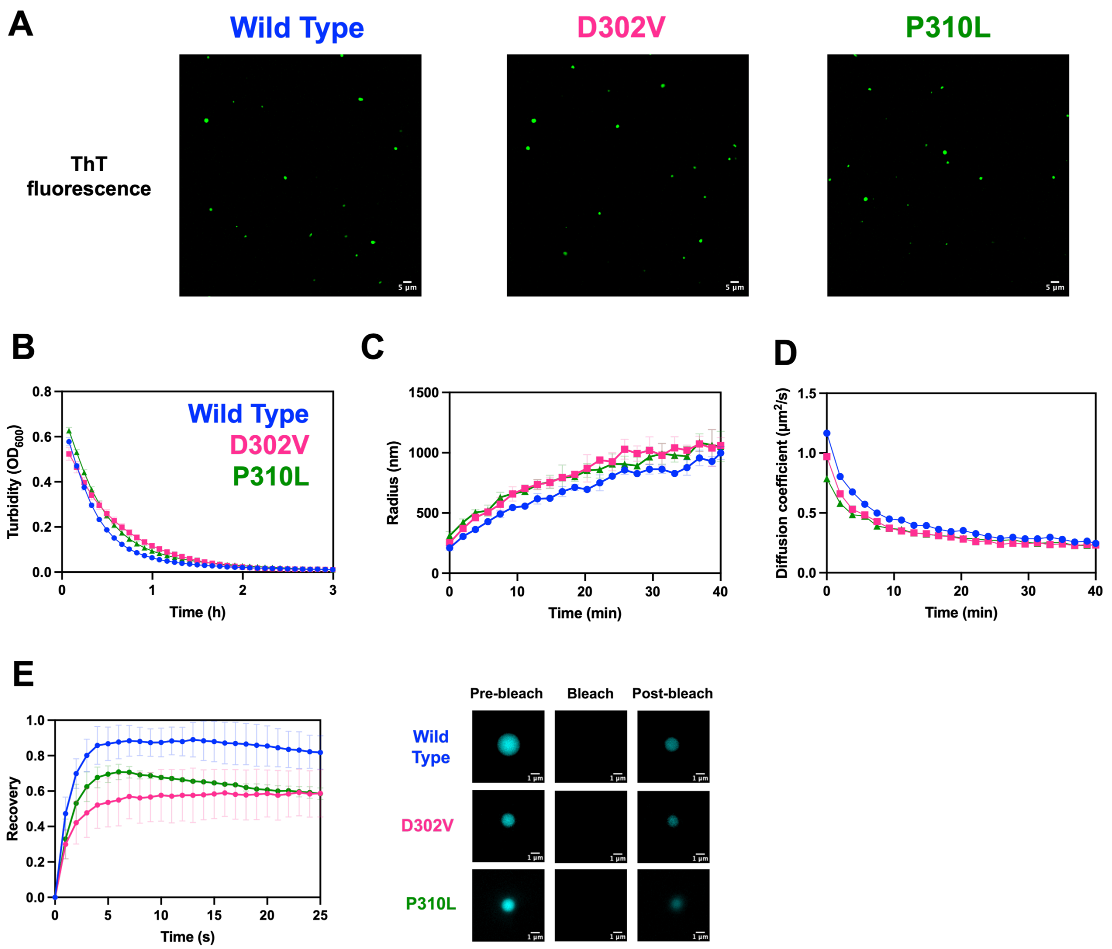


**Fig. S1. Biophysical characterization of phase separation and condensate dynamics of hnRNPA2B1 LCD variants**

**A)** Representative confocal thioflavin T (ThT) fluorescence microscopy of WT, D302V, and 310L hnRNPA2B1 LCD variants, showing ThT-positive condensates. Scale bar: 5 µm. **B)** LLPS kinetics monitored by turbidity ($OD_{600}$) measurements. Data are presented as the mean ± SD of three technical replicates. **C)** Average hydrodynamic radius and **D)** diffusion coefficients of WT, D302V, and P310L condensates over time, determined by DLS. Data are presented as the mean ± SD of three independent experiments. **E)** FRAP recovery of WT, D302V, and P310L condensates with representative pre-bleach, bleach, and post-bleach images. Data are presented as the mean ± SD of five condensates. Scale bars: 1 µm.

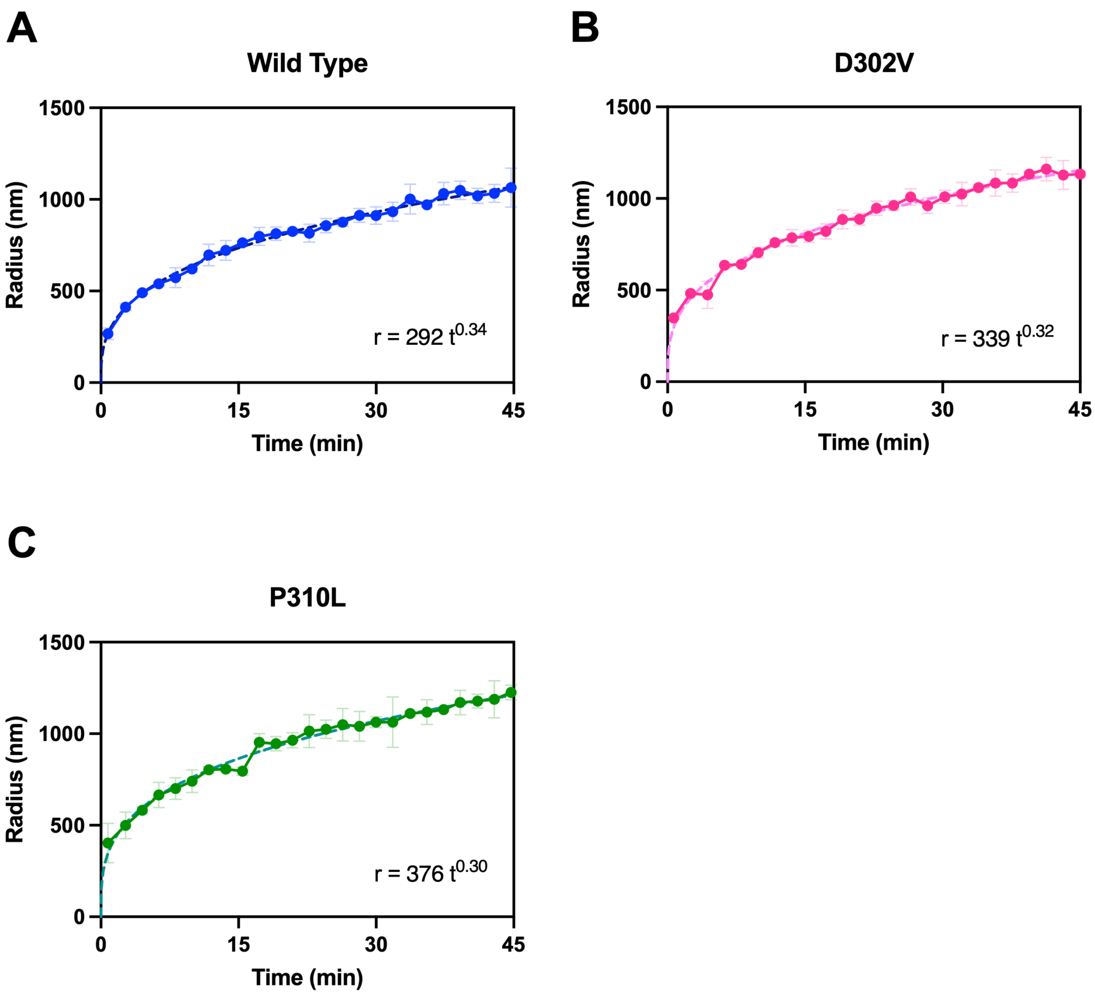


**Fig. S2**. **Droplet growth kinetics of full-length hnRNPA2B1 variants**
Average hydrodynamic radius of **A)** wild-type, **B)** D302V, and **C)** P310L hnRNPA2B1 condensates measured by dynamic Light Scattering (DLS). Condensates growth follows an approximately $t^{1/3}$ scaling law, consistent with diffusion-governed coarsening, through mechanisms such as Ostwald ripening and collision-induced coalescence. Dotted lines represent the fits to the indicated power-law equations. Data are presented as the mean ± SD of three independent experiments.

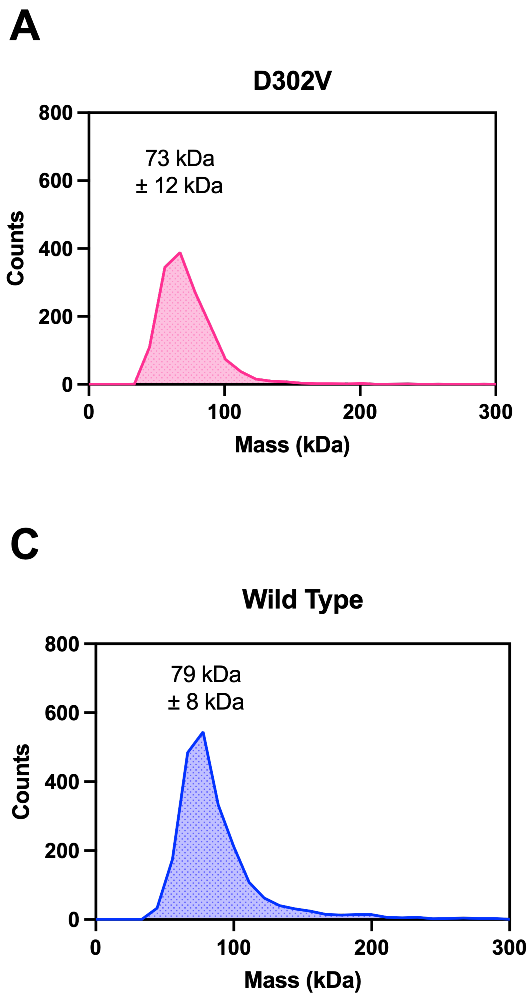


**Fig. S3**. **Disease-linked mutations do not alter the oligomeric state of hnRNPA2B1**
Mass photometry analysis of **A)** D302V and **B)** P310L hnRNPA2B1 mutants determined apparent molecular masses of 73 ± 12 kDa and 68 ± 2 kDa, respectively. **C)** Wild type hnRNPA2B1 shows an apparent molecular mass of 79 ± 8 kDa and is included as a reference from our previous publication (1). Distributions represent the detected particle counts as a function of molecular mass. Each variant was measured in triplicate. Values indicate the mean apparent molecular mass ± SD across the three measurements.

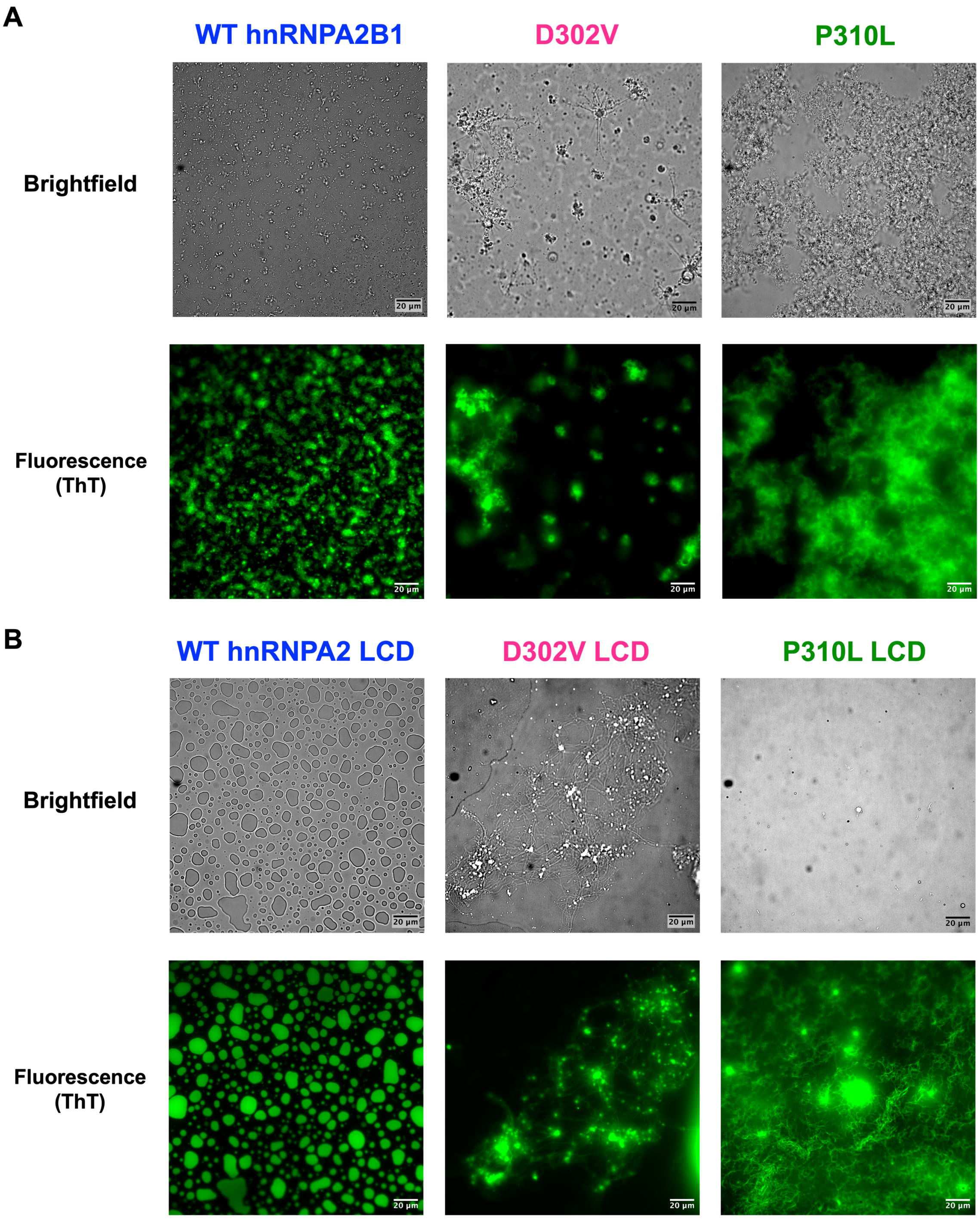


**Fig. S4. Morphology of ThT-positive structures formed by full-length hnRNPA2B1 and isolated LCD variants.** Representative brightfield and ThT fluorescence microscopy images of **A)** WT, D302V, and P310L full-length hnRNPA2B1 and **B)** WT, D302V, and P310L hnRNPA2B1 LCD following overnight incubation after induction of phase separation. ThT fluorescence confirms the presence of β-sheet-rich structures in both full-length and isolated LCD constructs. Scale bars: 20 µm.

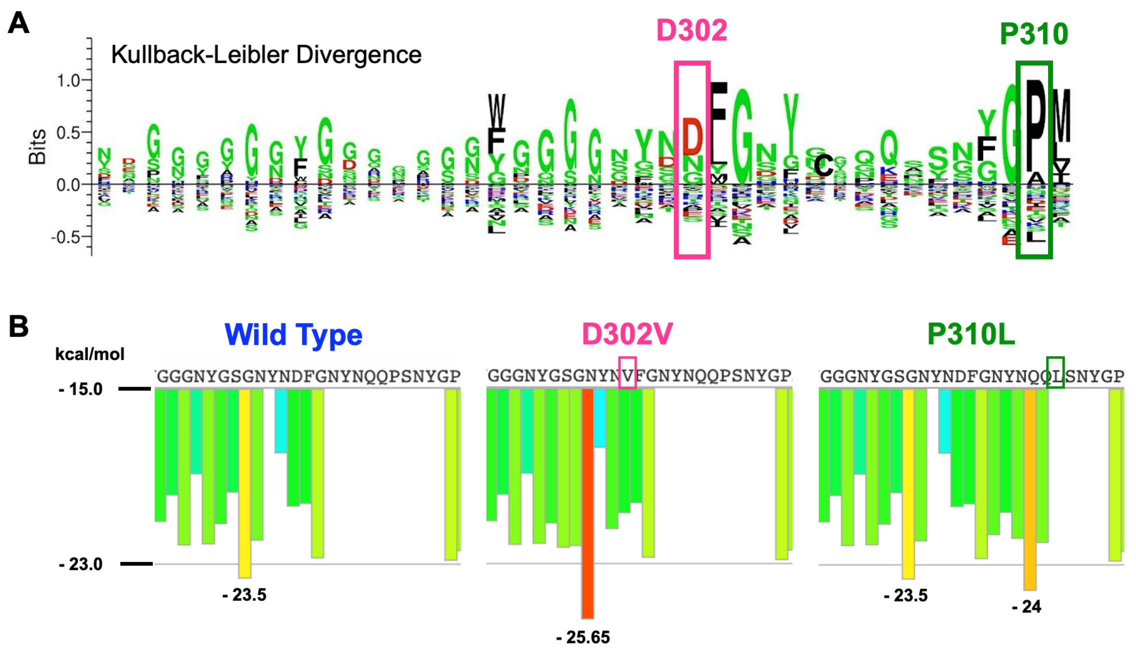


**Fig. S5**. **Disease-linked mutations reside within an evolutionarily conserved region within the LCD of hNRPA2B1**

**A)** Kullback-Leibler (KL) sequence logo generated from a MSA of 392 vertebrate hnRNPA2B1 LCD orthologs. Letter height reflects the degree of enrichment or depletion of each amino acid at a position (large = highly enriched or depleted). Letters above the x-axis represent evolutionarily conserved residues at that position, ordered from most to least frequent (top to bottom). Letters below the x-axis indicate residues that are disfavored, ordered from less to most depleted (top to bottom). At position 302 (magenta box), D is enriched, with N and G appearing as mildly tolerated alternative, consistent with their overall enrichment along the LCD and in the steric zipper motif. At position 310 (green box), P is extremely conserved. Both pathogenic mutations (V302 and L310) are strongly disfavored at their corresponding positions, highlighting strong evolutionary pressure against introducing hydrophobicity. **B)** ZipperDB predictions of the amyloidogenic propensity of hexapeptides in the region between G291-P315 of the LCD of hnRNPA2B1. The native sequence harbors a steric zipper motif GNYNDF, which is drives amyloid formation (yellow bar). The D302V mutation increases the strength of the native steric zipper (red bar), lowering the Rosetta energy for fibrillization. The P310L mutation introduces a new amyloidogenic hexapeptide QQLSNY just four residues adjacent of the native steric zipper (orange bar).

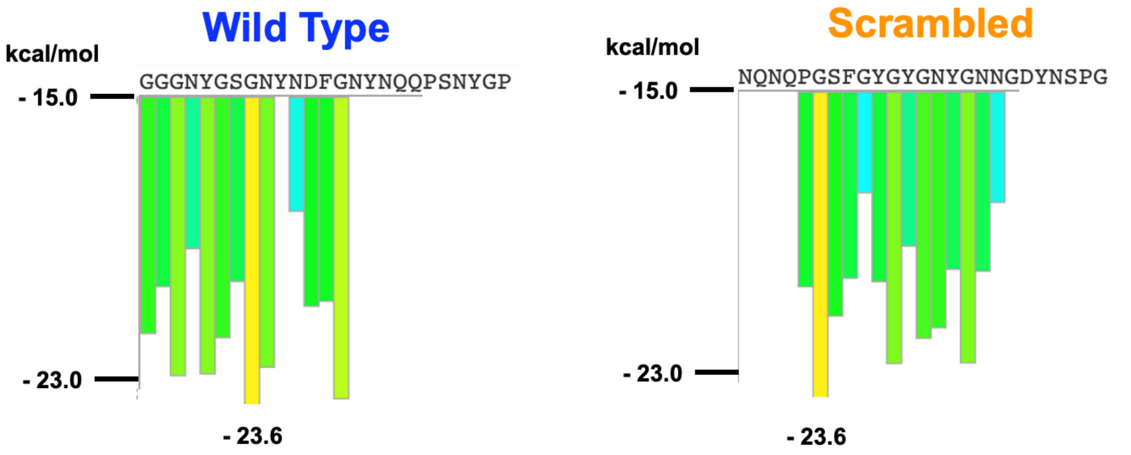


**Fig. S6. ZipperDB analysis of WT and Scrambled Amy peptides**
The selected scrambled sequence disrupts the native steric zipper motif while retaining a predicted amyloidogenic profile comparable to the WT peptide.

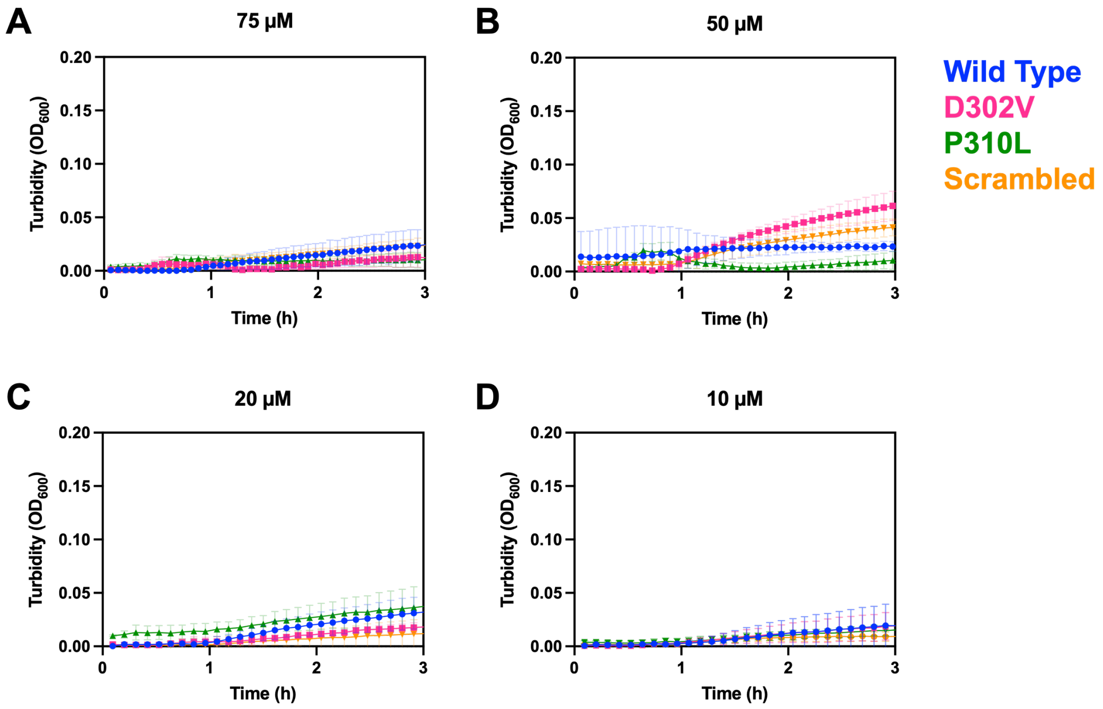


**Fig. S7. Amy peptides do not undergo detectable phase separation**
Turbidity measurements at 600 nm of wild-type (blue), D302V (pink), P310L (green), and scrambled (orange) Amy peptides at different concentrations: **A)** 75 μM, **B)** 50 μM, **C)** 20 μM, and **D)** 10 μM. Across all conditions, the turbidity remained at the baseline, indicating absence of phase separation. Data are presented as the mean ± SD of three technical replicates.

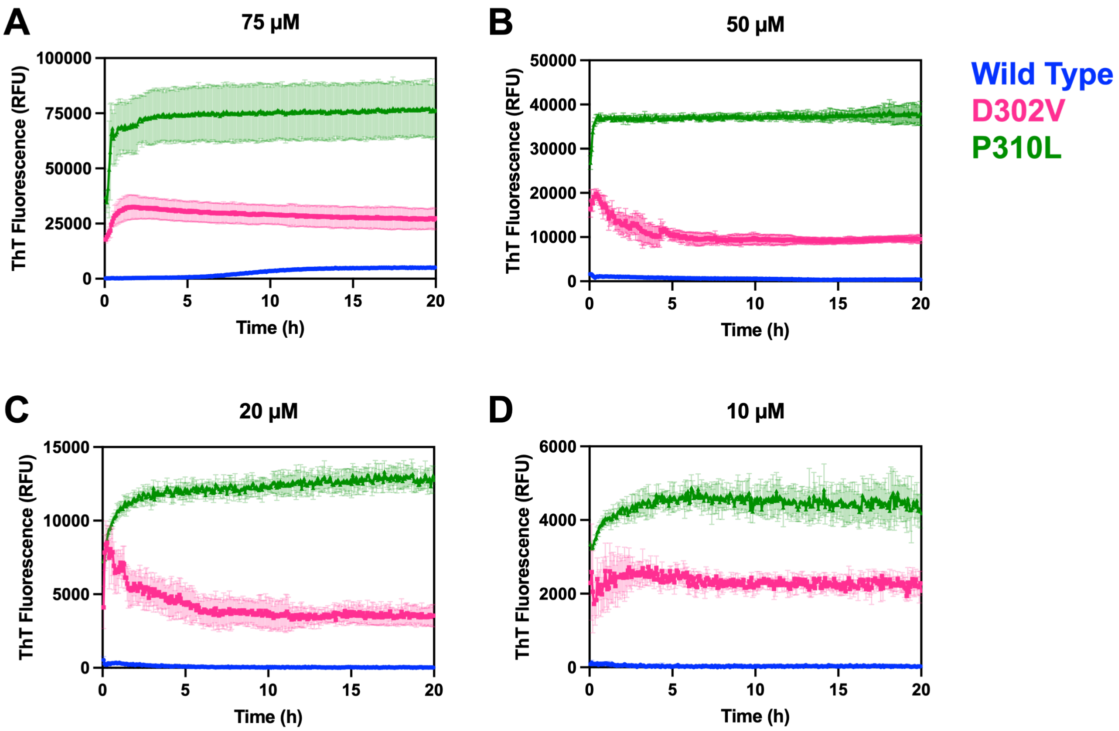


**Fig. S8. Mutant Amy peptides display enhanced aggregation propensity across different concentrations.**

ThT fluorescence kinetics of wild-type (blue), D302V (pink), and P310L (green) Amy peptides at different concentrations. **A)** At 75 µM, all Amy peptides follow sigmoidal growth kinetics, characteristic of amyloid-like aggregation. Both mutants exhibit faster aggregation kinetics with enhanced ThT fluorescence, being P310L the strongest. **B)** At 50 µM, the wild-type peptide remains at baseline. However, the D302V and P310L mutants still display high aggregation propensity. This trend is also observed at **C)** 20 µM and **D)** 10 µM, suggesting that the mutations increase the aggregation propensity, even below the critical concentration for aggregation of the wild-type peptide. Data are presented as the mean ± SD of three technical replicates.

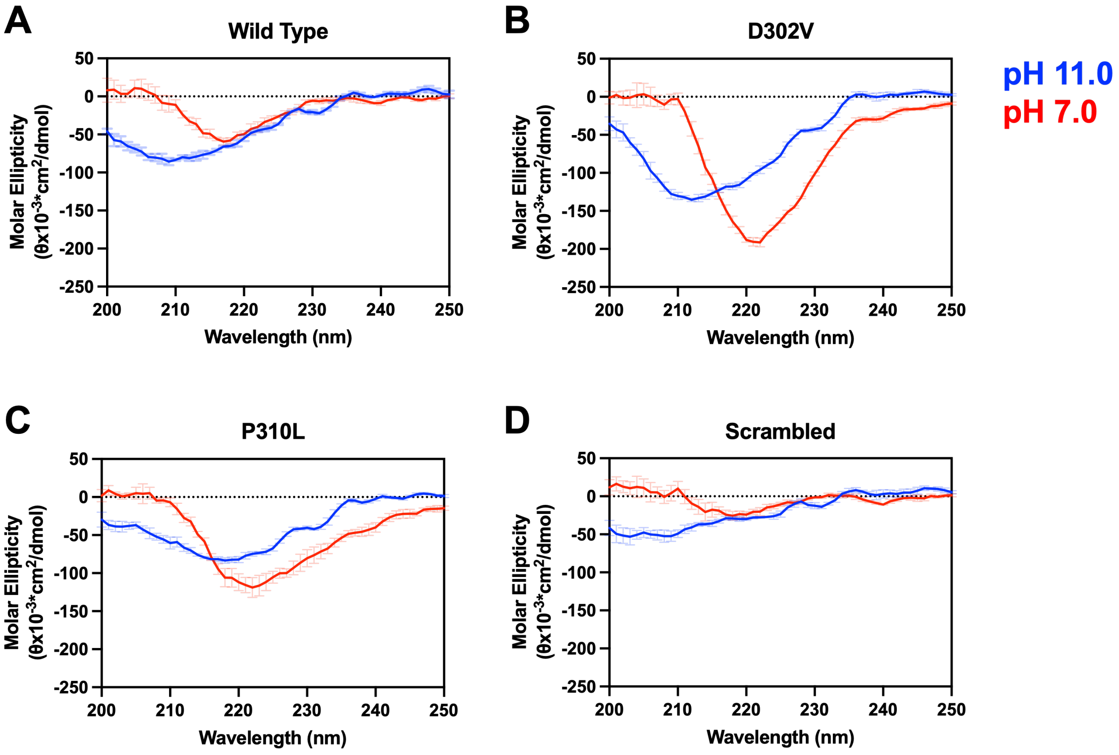


**Fig. S9. The Amy peptides undergo pH-dependent conformational changes**
Circular dichroism spectra of **A)** WT, **B)** D302V, **C)** P310L, and **D)** scrambled Amy peptides recorded in storage buffer (10 mM CAPS, pH 11.0; blue) and upon dilution into physiological pH (50 mM HEPES, pH 7.0; red). Although the Amy peptides remain soluble at high pH, WT and mutant peptides retain residual β-sheet propensity. Following the pH jump, they exhibited a shift in ellipticity toward 218-220 nm, indicative of β-sheet-rich conformations. The scrambled control remained predominantly disordered in both conditions. Data are presented as the mean ± SD of three independent experiments.

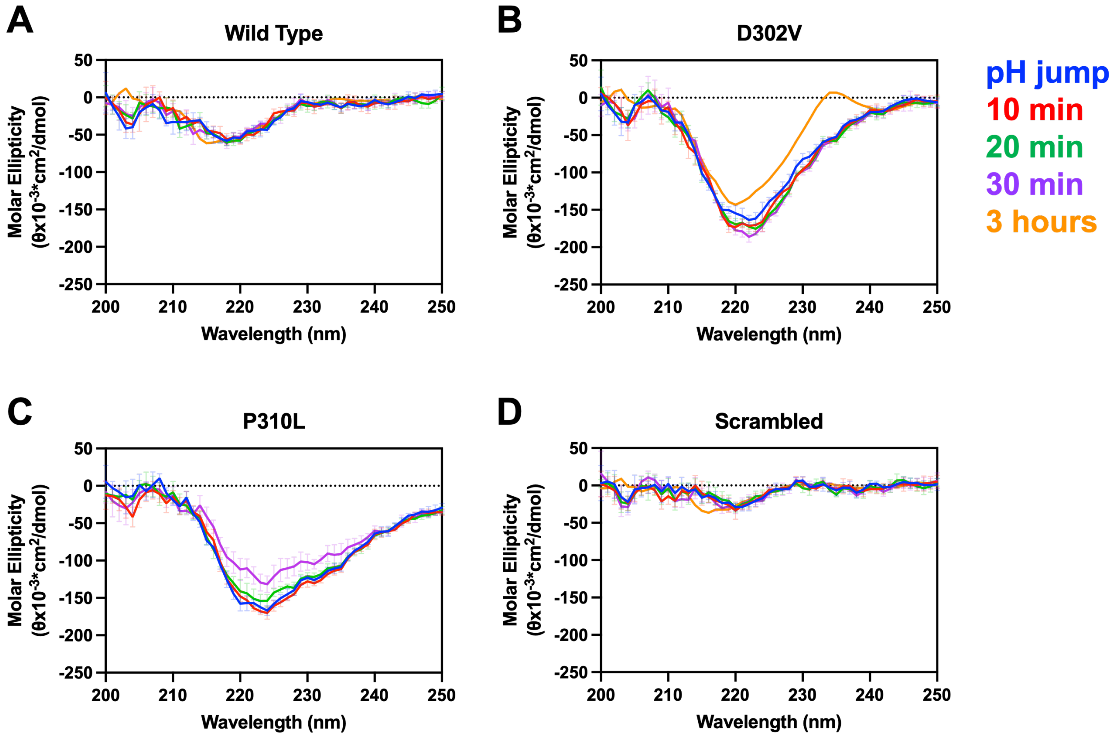


**Fig. S10. Timelapse of secondary structure analysis of the Amy peptides by CD spectroscopy**

CD spectra of Amy peptides at different timepoints. Spectra were recorded immediately upon pH jump (time 0; blue) and after 10 (red), 20 (green), 30 (purple) minutes, and 3 hours (yellow) of incubation. **A)** WT peptide exhibits a moderate β-sheet-like spectra, with minimums around 218-220 nm. This signature remained stable throughout the 3 hours incubation, indicating a slower aggregation pathway. **B)** D302V displays a strong signal for β-sheet content, slightly increasing over the first 30 minutes, and reducing after 3 hours due to slow aggregation. **C)** P310L aggregates the fastest, showing immediate β-sheet formation and aggregation in the cuvette within 30 minutes of incubation. **D)** The scrambled control remains largely unstructured and does not undergo any major structural ordering, reinforcing its non-aggregating nature. Data from 0 to 30 minutes represent the mean of three independent experiments, whereas the 3h spectra are representative of the mean of technical replicates.

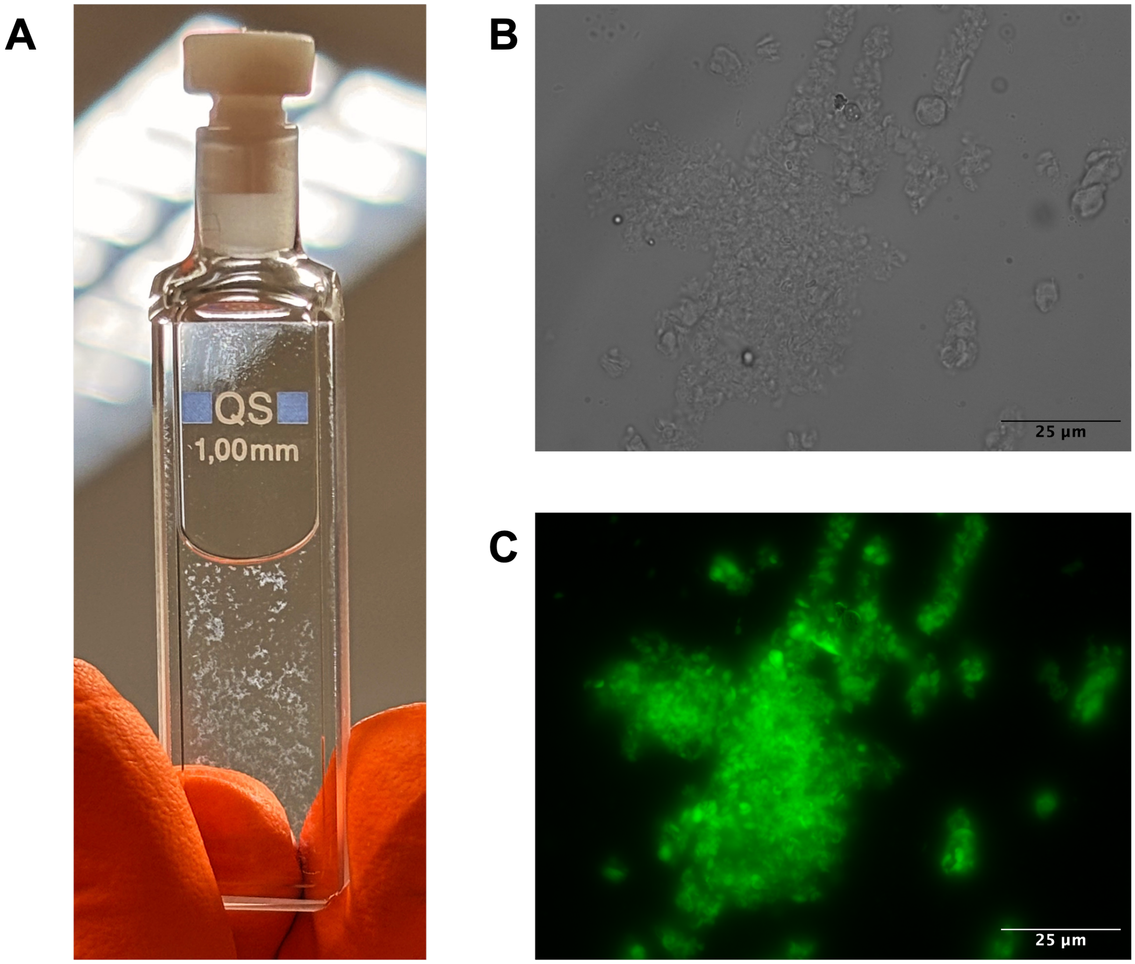


**Fig. S11.The P310L peptide aggregates during CD measurements**
**A)** Aggregates could be observed in the cuvette during CD measurements, even at early timepoints. **B)** Brightfield and **C)** Fluorescence microscopy of P310L peptide stained with ThT. Scale bar: 25 µm.

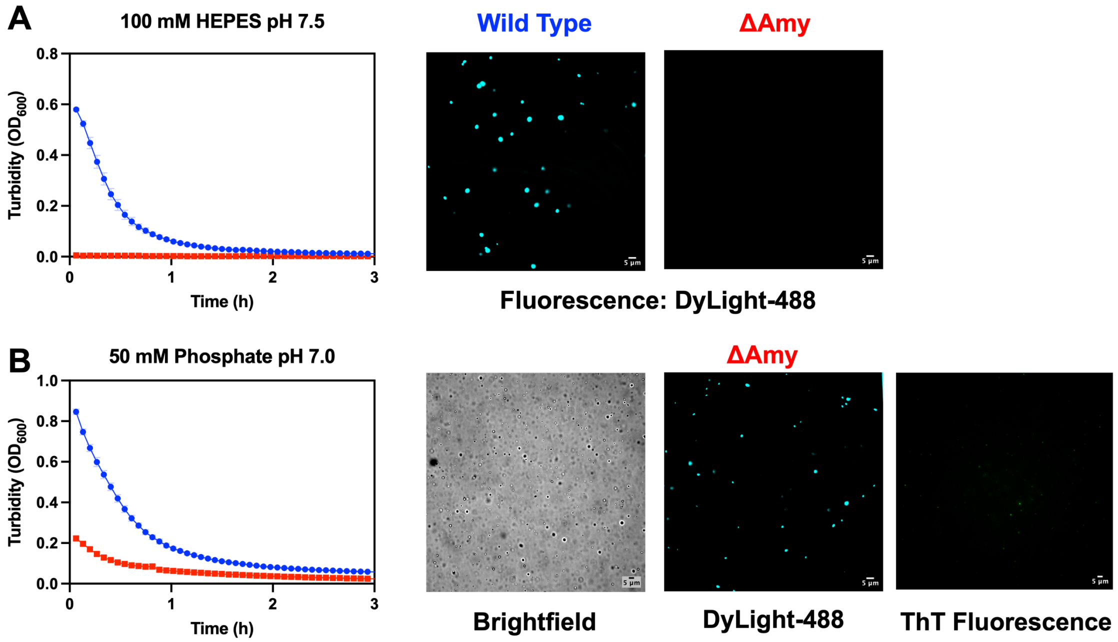


**Fig. S12. Deletion of the amyloidogenic region severely disrupts phase separation behavior. A)** LLPS of WT hnRNPA2 LCD (blue) and ΔAmy (red) in 100 mM HEPES, pH 7.5, monitored by turbidity measurements ($OD_{600}$) and fluorescence microscopy with DyLight488™ NHS Ester-labeled protein. ΔAmy does not exhibit detectable phase separation under these conditions. **B)** LLPS of WT and ΔAmy in 50 mM phosphate buffer, pH 7.0. Under these conditions, ΔAmy undergoes LLPS, although with a heavily reduced propensity when compared to wild-type protein. Confocal fluorescence microscopy shows that ΔAmy condensates exhibit little to no binding to ThT. Data are presented as the mean ± SD of three technical replicates. Scale bars: 5 µm.

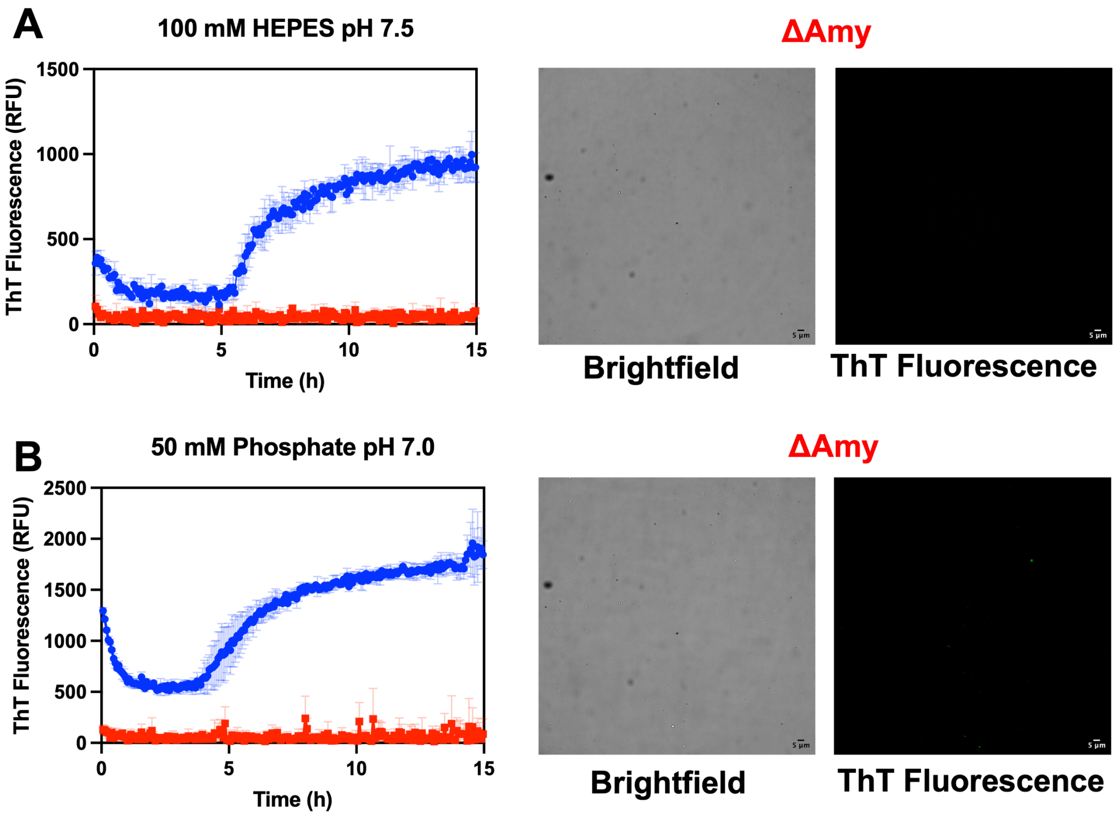


**Fig.S13. Deletion of the Amy region abolishes amyloid-like aggregation of hnRNPA2 LCD**
ThT fluorescence kinetics of wild type hnRNPA2 LCD (blue) and ΔAmy (red) hnRNPA2 LCD solutions in **A)** 100 mM HEPES pH 7.5 and **B)** 50 mM phosphate pH 7.0. Under both conditions, wild type hnRNPA2 LCD follows sigmoidal ThT fluorescence kinetics, whereas ΔAmy signal remains near baseline throughout the measurement. Corresponding brightfield and ThT fluorescence microscopy show no detectable structures for ΔAmy, confirming that deletion of the Amy region abolishes detectable amyloid-like aggregation under these conditions. Data are presented as the mean ± SD of three technical replicates. Scale bars: 5 μm.

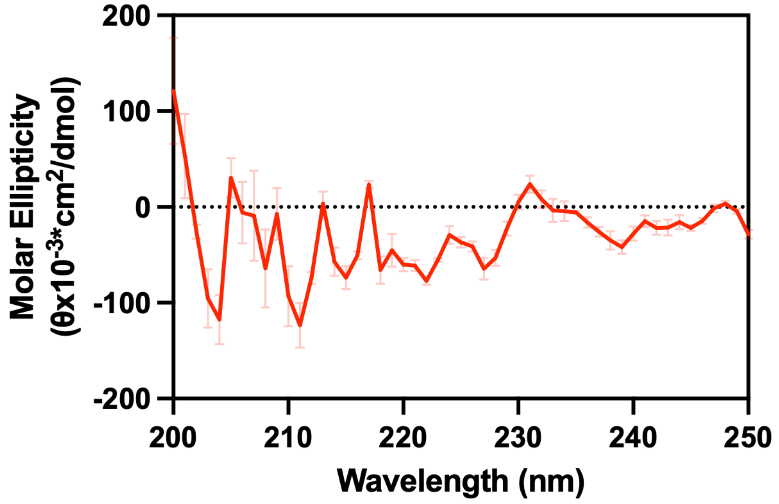


**Fig.S14. ΔAmy does not exhibit β-sheet-rich conformations.**
Circular dichroism (CD) spectrum of ΔAmy hnRNPA2 LCD in 50 mM HEPES, pH 7.0. Contrary to the Amy peptides, ΔAmy does not exhibit a clearly defined β-sheet-like spectral signature under these conditions, consistent with the lack of ThT fluorescence and amyloid-like aggregation. Data represent the mean ± SD of three consecutive measurements.

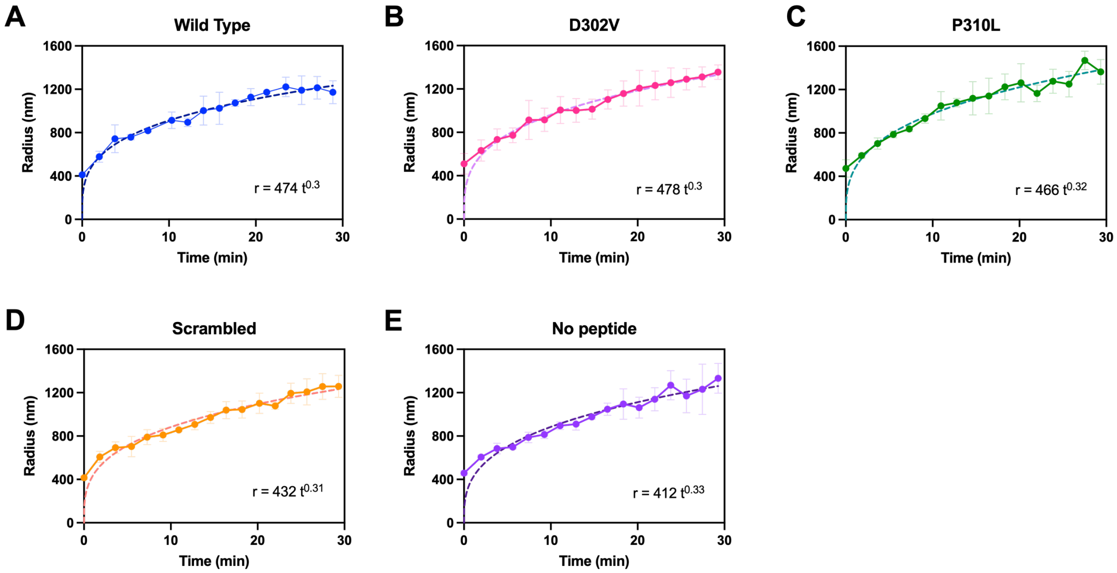


**Fig. S15**. **Droplet growth kinetics of hnRNPA2 LCD seeded with Amy fibrils**
Average hydrodynamic radius of hnRNPA2 LCD seeded with pre-formed **A)** wild-type, **B)** D302V, **C)** P310L, and **D)** scrambled fibrils compared to the **E)** unseeded control. Addition of pre-formed fibrils does not markedly accelerate condensate growth kinetics compared to the unseeded control, with all conditions exhibiting $t^{1/3}$ scaling, consistent with coarsening. Dotted lines represent the fits to the indicated power-law equations. Data are presented as the mean ± SD of three independent experiments.

### Tables

#### Table S1. ZipperDB scores (kcal/mol) for predicted amyloidogenic regions in the LCD of hnRNPA2B1.

| | Wild Type | | D302V | | P310L | |
|---|---|---|---|---|---|---|
| **Position** | **Hexapeptide** | **Score** | **Hexapeptide** | **Score** | **Hexapeptide** | **Score** |
| 1 | RSGRGG | -10.68342 | RSGRGG | -10.68342 | RSGRGG | -10.68342 |
| 2 | SGRGGN | -15.47103 | SGRGGN | -15.47103 | SGRGGN | -15.47103 |
| 3 | GRGGNF | -17.90026 | GRGGNF | -17.90026 | GRGGNF | -17.90026 |
| 4 | RGGNFG | -18.53729 | RGGNFG | -18.53729 | RGGNFG | -18.53729 |
| 5 | GGNFGF | -19.45553 | GGNFGF | -19.45553 | GGNFGF | -19.45553 |
| 6 | GNFGFG | -22.8207 | GNFGFG | -22.8207 | GNFGFG | -22.8207 |
| 7 | NFGFGD | -15.78282 | NFGFGD | -15.78282 | NFGFGD | -15.78282 |
| 8 | FGFGDS | -18.55468 | FGFGDS | -18.55468 | FGFGDS | -18.55468 |
| 9 | GFGDSR | -14.63815 | GFGDSR | -14.63815 | GFGDSR | -14.63815 |
| 10 | FGDSRG | -14.17159 | FGDSRG | -14.17159 | FGDSRG | -14.17159 |
| 11 | GDSRGG | -16.60894 | GDSRGG | -16.60894 | GDSRGG | -16.60894 |
| 12 | DSRGGG | -15.21133 | DSRGGG | -15.21133 | DSRGGG | -15.21133 |
| 13 | SRGGGG | -14.79151 | SRGGGG | -14.79151 | SRGGGG | -14.79151 |
| 14 | RGGGGN | -12.94155 | RGGGGN | -12.94155 | RGGGGN | -12.94155 |
| 15 | GGGGNF | -21.8887 | GGGGNF | -21.8887 | GGGGNF | -21.8887 |
| 16 | GGGNFG | -21.73927 | GGGNFG | -21.73927 | GGGNFG | -21.73927 |
| 17 | GGNFGP | NaN | GGNFGP | NaN | GGNFGP | NaN |
| 18 | GNFGPG | 49.07635 | GNFGPG | 49.07635 | GNFGPG | 49.07635 |
| 19 | NFGPGP | NaN | NFGPGP | NaN | NFGPGP | NaN |
| 20 | FGPGPG | 64.04657 | FGPGPG | 64.04657 | FGPGPG | 64.04657 |
| 21 | GPGPGS | 68.35456 | GPGPGS | 68.35456 | GPGPGS | 68.35456 |
| 22 | PGPGSN | NaN | PGPGSN | NaN | PGPGSN | NaN |
| 23 | GPGSNF | 57.20353 | GPGSNF | 57.20353 | GPGSNF | 57.20353 |
| 24 | PGSNFR | NaN | PGSNFR | NaN | PGSNFR | NaN |
| 25 | GSNFRG | -16.06997 | GSNFRG | -16.06997 | GSNFRG | -16.06997 |
| 26 | SNFRGG | -15.04685 | SNFRGG | -15.04685 | SNFRGG | -15.04685 |
| 27 | NFRGGS | -13.62501 | NFRGGS | -13.62501 | NFRGGS | -13.62501 |
| 28 | FRGGSD | -12.21711 | FRGGSD | -12.21711 | FRGGSD | -12.21711 |
| 29 | RGGSDG | -14.58421 | RGGSDG | -14.58421 | RGGSDG | -14.58421 |
| 30 | GGSDGY | -21.75274 | GGSDGY | -21.75274 | GGSDGY | -21.75274 |
| 31 | GSDGYG | -20.37394 | GSDGYG | -20.37394 | GSDGYG | -20.37394 |
| 32 | SDGYGS | -16.4797 | SDGYGS | -16.4797 | SDGYGS | -16.4797 |
| 33 | DGYGSG | -20.05544 | DGYGSG | -20.05544 | DGYGSG | -20.05544 |
| 34 | GYGSGR | -16.26356 | GYGSGR | -16.26356 | GYGSGR | -16.26356 |
| 35 | YGSGRG | -14.73938 | YGSGRG | -14.73938 | YGSGRG | -14.73938 |
| 36 | GSGRGF | -17.59933 | GSGRGF | -17.59933 | GSGRGF | -17.59933 |
| 37 | SGRGFG | -16.903 | SGRGFG | -16.903 | SGRGFG | -16.903 |
| 38 | GRGFGD | -11.92437 | GRGFGD | -11.92437 | GRGFGD | -11.92437 |
| 39 | RGFGDG | -15.89512 | RGFGDG | -15.89512 | RGFGDG | -15.89512 |
| 40 | GFGDGY | -19.21158 | GFGDGY | -19.21158 | GFGDGY | -19.21158 |
| 41 | FGDGYN | -17.122 | FGDGYN | -17.122 | FGDGYN | -17.122 |
| 42 | GDGYNG | -18.12914 | GDGYNG | -18.12914 | GDGYNG | -18.12914 |
| 43 | DGYNGY | -20.10479 | DGYNGY | -20.10479 | DGYNGY | -20.10479 |
| 44 | GYNGYG | -24.22609 | GYNGYG | -24.22609 | GYNGYG | -24.22609 |
| 45 | YNGYGG | -16.81361 | YNGYGG | -16.81361 | YNGYGG | -16.81361 |
| 46 | NGYGGG | -17.60922 | NGYGGG | -17.60922 | NGYGGG | -17.60922 |
| 47 | GYGGGP | NaN | GYGGGP | NaN | GYGGGP | NaN |
| 48 | YGGGPG | 28.67514 | YGGGPG | 28.67514 | YGGGPG | 28.67514 |
| 49 | GGGPGG | -6.54477 | GGGPGG | -6.54477 | GGGPGG | -6.54477 |
| 50 | GGPGGG | 1.1434 | GGPGGG | 1.1434 | GGPGGG | 1.1434 |
| 51 | GPGGGN | 70.79393 | GPGGGN | 70.79393 | GPGGGN | 70.79393 |
| 52 | PGGGNF | NaN | PGGGNF | NaN | PGGGNF | NaN |
| 53 | GGGNFG | -21.73927 | GGGNFG | -21.73927 | GGGNFG | -21.73927 |
| 54 | GGNFGG | -18.90319 | GGNFGG | -18.90319 | GGNFGG | -18.90319 |
| 55 | GNFGGS | -21.39888 | GNFGGS | -21.39888 | GNFGGS | -21.39888 |
| 56 | NFGGSP | NaN | NFGGSP | NaN | NFGGSP | NaN |
| 57 | FGGSPG | 14.67633 | FGGSPG | 14.67633 | FGGSPG | 14.67633 |
| 58 | GGSPGY | -10.0839 | GGSPGY | -10.0839 | GGSPGY | -10.0839 |
| 59 | GSPGYG | 2.43509 | GSPGYG | 2.43509 | GSPGYG | 2.43509 |
| 60 | SPGYGG | 66.91133 | SPGYGG | 66.91133 | SPGYGG | 66.91133 |
| 61 | PGYGGG | NaN | PGYGGG | NaN | PGYGGG | NaN |
| 62 | GYGGGR | -14.82642 | GYGGGR | -14.82642 | GYGGGR | -14.82642 |
| 63 | YGGGRG | -13.8923 | YGGGRG | -13.8923 | YGGGRG | -13.8923 |
| 64 | GGGRGG | -14.29839 | GGGRGG | -14.29839 | GGGRGG | -14.29839 |
| 65 | GGRGGY | -15.37504 | GGRGGY | -15.37504 | GGRGGY | -15.37504 |
| 66 | GRGGYG | -16.53343 | GRGGYG | -16.53343 | GRGGYG | -16.53343 |
| 67 | RGGYGG | -12.9683 | RGGYGG | -12.9683 | RGGYGG | -12.9683 |
| 68 | GGYGGG | -20.27983 | GGYGGG | -20.27983 | GGYGGG | -20.27983 |
| 69 | GYGGGG | -19.11935 | GYGGGG | -19.11935 | GYGGGG | -19.11935 |
| 70 | YGGGGP | NaN | YGGGGP | NaN | YGGGGP | NaN |
| 71 | GGGGPG | 25.99276 | GGGGPG | 25.99276 | GGGGPG | 25.99276 |
| 72 | GGGPGY | -8.14372 | GGGPGY | -8.14372 | GGGPGY | -8.14372 |
| 73 | GGPGYG | 3.14905 | GGPGYG | 3.14905 | GGPGYG | 3.14905 |

| | | | | | | |
|---|---|---|---|---|---|---|
| 74 | GPGYGN | 70.26051 | GPGYGN | 70.26051 | GPGYGN | 70.26051 |
| 75 | PGYGNQ | NaN | PGYGNQ | NaN | PGYGNQ | NaN |
| 76 | GYGNQG | -21.74202 | GYGNQG | -21.74202 | GYGNQG | -21.74202 |
| 77 | YGNQGG | -21.03761 | YGNQGG | -21.03761 | YGNQGG | -21.03761 |
| 78 | GNQGGG | -20.39547 | GNQGGG | -20.39547 | GNQGGG | -20.39547 |
| 79 | NQGGGY | -19.85661 | NQGGGY | -19.85661 | NQGGGY | -19.85661 |
| 80 | QGGGYG | -22.18077 | QGGGYG | -22.18077 | QGGGYG | -22.18077 |
| 81 | GGGYGG | -17.4265 | GGGYGG | -17.4265 | GGGYGG | -17.4265 |
| 82 | GGYGGG | -20.27983 | GGYGGG | -20.27983 | GGYGGG | -20.27983 |
| 83 | GYGGGY | -20.41021 | GYGGGY | -20.41021 | GYGGGY | -20.41021 |
| 84 | YGGGYD | -20.00542 | YGGGYD | -20.00542 | YGGGYD | -20.00542 |
| 85 | GGGYDN | -15.51213 | GGGYDN | -15.51213 | GGGYDN | -15.51213 |
| 86 | GGYDNY | -22.55827 | GGYDNY | -22.55827 | GGYDNY | -22.55827 |
| 87 | GYDNYG | -20.8994 | GYDNYG | -20.8994 | GYDNYG | -20.8994 |
| 88 | YDNYGG | -17.68269 | YDNYGG | -17.68269 | YDNYGG | -17.68269 |
| 89 | DNYGGG | -18.00789 | DNYGGG | -18.00789 | DNYGGG | -18.00789 |
| 90 | NYGGGN | -17.81208 | NYGGGN | -17.81208 | NYGGGN | -17.81208 |
| 91 | YGGGNY | -20.86494 | YGGGNY | -20.86494 | YGGGNY | -20.86494 |
| 92 | GGGNYG | -21.1801 | GGGNYG | -21.1801 | GGGNYG | -21.1801 |
| 93 | GGNYGS | -19.95688 | GGNYGS | -19.95688 | GGNYGS | -19.95688 |
| 94 | GNYGSG | -22.04076 | GNYGSG | -22.04076 | GNYGSG | -22.04076 |
| 95 | NYGSGN | -18.74706 | NYGSGN | -18.74706 | NYGSGN | -18.74706 |
| 96 | YGSGNY | -21.82486 | YGSGNY | -21.82486 | YGSGNY | -21.82486 |
| 97 | GSGNYN | -21.3253 | GSGNYN | -21.3253 | GSGNYN | -21.3253 |
| **98** | SGNYND | -19.8379 | **SGNYNV** | **-22.4504** | SGNYND | -19.8379 |
| **99** | **GNYNDF** | **-23.4921** | **GNYNVF** | **-22.14393** | **GNYNDF** | **-23.4921** |
| **100** | NYNDFG | -21.96992 | **NYNVFG** | **-25.65154** | NYNDFG | -21.96992 |
| **101** | YNDFGN | -13.69056 | **YNVFGN** | **-17.76783** | YNDFGN | -13.69056 |
| **102** | NDFGNY | -18.34871 | **NVFGNY** | **-21.71734** | NDFGNY | -18.34871 |
| **103** | DFGNYN | -20.45489 | **VFGNYN** | **-20.9594** | DFGNYN | -20.45489 |
| **104** | FGNYNQ | -20.05463 | FGNYNQ | -20.05463 | FGNYNQ | -20.05463 |
| **105** | GNYNQQ | -22.8164 | GNYNQQ | -22.8164 | GNYNQQ | -22.8164 |
| **106** | NYNQQP | NaN | NYNQQP | NaN | **NYNQQL** | **-21.2844** |
| **107** | YNQQPS | 34.34046 | YNQQPS | 34.34046 | **YNQQLS** | **-20.61998** |
| **108** | NQQPSN | -6.73046 | NQQPSN | -6.73046 | **NQQLSN** | **-21.63088** |
| **109** | QQPSNY | 0.34435 | QQPSNY | 0.34435 | **QQLSNY** | **-23.93095** |
| **110** | QPSNYG | 57.81308 | QPSNYG | 57.81308 | **QLSNYG** | **-21.97004** |
| 111 | PSNYGP | NaN | PSNYGP | NaN | LSNYGP | NaN |
| 112 | SNYGPM | 38.21611 | SNYGPM | 38.21611 | SNYGPM | 38.21611 |
| 113 | NYGPMK | -8.0629 | NYGPMK | -8.0629 | NYGPMK | -8.0629 |
| 114 | YGPMKS | 9.79121 | YGPMKS | 9.79121 | YGPMKS | 9.79121 |
| 115 | GPMKSG | 64.83423 | GPMKSG | 64.83423 | GPMKSG | 64.83423 |
| 116 | PMKSGN | NaN | PMKSGN | NaN | PMKSGN | NaN |
| 117 | MKSGNF | -22.55485 | MKSGNF | -22.55485 | MKSGNF | -22.55485 |
| 118 | KSGNFG | -22.5836 | KSGNFG | -22.5836 | KSGNFG | -22.5836 |
| 119 | SGNFGG | -18.98122 | SGNFGG | -18.98122 | SGNFGG | -18.98122 |
| 120 | GNFGGS | -21.39888 | GNFGGS | -21.39888 | GNFGGS | -21.39888 |
| 121 | NFGGSR | -15.38136 | NFGGSR | -15.38136 | NFGGSR | -15.38136 |
| 122 | FGGSRN | -12.14021 | FGGSRN | -12.14021 | FGGSRN | -12.14021 |
| 123 | GGSRNM | -18.63083 | GGSRNM | -18.63083 | GGSRNM | -18.63083 |
| 124 | GSRNMG | -16.49164 | GSRNMG | -16.49164 | GSRNMG | -16.49164 |
| 125 | SRNMGG | -17.06463 | SRNMGG | -17.06463 | SRNMGG | -17.06463 |
| 126 | RNMGGP | NaN | RNMGGP | NaN | RNMGGP | NaN |
| 127 | NMGGPY | 21.59679 | NMGGPY | 21.59679 | NMGGPY | 21.59679 |
| 128 | MGGPYG | -10.40851 | MGGPYG | -10.40851 | MGGPYG | -10.40851 |
| 129 | GGPYGG | 1.00834 | GGPYGG | 1.00834 | GGPYGG | 1.00834 |
| 130 | GPYGGG | 62.41519 | GPYGGG | 62.41519 | GPYGGG | 62.41519 |
| 131 | PYGGGN | NaN | PYGGGN | NaN | PYGGGN | NaN |
| 132 | YGGGNY | -20.86494 | YGGGNY | -20.86494 | YGGGNY | -20.86494 |
| 133 | GGGNYG | -21.1801 | GGGNYG | -21.1801 | GGGNYG | -21.1801 |
| 134 | GGNYGP | NaN | GGNYGP | NaN | GGNYGP | NaN |
| 135 | GNYGPG | 42.75663 | GNYGPG | 42.75663 | GNYGPG | 42.75663 |
| 136 | NYGPGG | -6.57471 | NYGPGG | -6.57471 | NYGPGG | -6.57471 |
| 137 | YGPGGS | 4.60132 | YGPGGS | 4.60132 | YGPGGS | 4.60132 |
| 138 | GPGGSG | 66.4521 | GPGGSG | 66.4521 | GPGGSG | 66.4521 |
| 139 | PGGSGG | NaN | PGGSGG | NaN | PGGSGG | NaN |
| 140 | GGSGGS | -21.28966 | GGSGGS | -21.28966 | GGSGGS | -21.28966 |
| 141 | GSGGSG | -19.73752 | GSGGSG | -19.73752 | GSGGSG | -19.73752 |
| 142 | SGGSGG | -19.63367 | SGGSGG | -19.63367 | SGGSGG | -19.63367 |
| 143 | GGSGGY | -22.29249 | GGSGGY | -22.29249 | GGSGGY | -22.29249 |
| 144 | GSGGYG | -21.55133 | GSGGYG | -21.55133 | GSGGYG | -21.55133 |
| 145 | SGGYGG | -17.92911 | SGGYGG | -17.92911 | SGGYGG | -17.92911 |
| 146 | GGYGGR | -16.00506 | GGYGGR | -16.00506 | GGYGGR | -16.00506 |
| 147 | GYGGRS | -15.9535 | GYGGRS | -15.9535 | GYGGRS | -15.9535 |
| 148 | YGGRSR | -9.53274 | YGGRSR | -9.53274 | YGGRSR | -9.53274 |
| 149 | GGRSRY | -12.69054 | GGRSRY | -12.69054 | GGRSRY | -12.69054 |
